# The activation of INF2 by Piezo1/Ca^2+^ is required for mesenchymal to amoeboid transition in confined environments

**DOI:** 10.1101/2023.06.23.546346

**Authors:** Neelakshi Kar, Alexa P. Caruso, Nicos Prokopiou, Jeremy S. Logue

**Affiliations:** Department of Regenerative and Cancer Cell Biology, Albany Medical College, 47 New Scotland Ave, Albany, NY 12208

**Keywords:** cancer, mesenchymal, amoeboid, microenvironment, bleb

## Abstract

To invade heterogenous tissues, transformed cells may undergo a mesenchymal to amoeboid transition (MAT). However, the molecular mechanisms regulating this transition are poorly defined. In invasive melanoma cells, we demonstrate that intracellular [Ca^2+^] increases with the degree of confinement in a Piezo1 dependent fashion. Moreover, Piezo1/Ca^2+^ is found to drive amoeboid and not mesenchymal migration in confined environments. Consistent with a model in which Piezo1 senses tension at the plasma membrane, the percentage of cells using amoeboid migration is further increased in undulating microchannels. Surprisingly, amoeboid migration was not promoted by myosin light chain kinase (MLCK), which is sensitive to intracellular [Ca^2+^]. Instead, we report that Piezo1/Ca^2+^ activates inverted formin-2 (INF2) to induce widespread actin cytoskeletal remodeling. Strikingly, the activation of INF2 is found to promote de-adhesion, which in turn facilitates MAT. Using micropatterned surfaces, we demonstrate that cells require INF2 to effectively migrate in environments with challenging mechanochemical properties.

**Summary Statement:** Migrating melanoma cells are found to rely on the activation of inverted formin-2 (INF2) by Piezo1/Ca^2+^ for mesenchymal to amoeboid transition (MAT) in confined environments.

## Introduction

To migrate within tissues, cells must adapt to diverse mechanochemical environments. For instance, immune cells must move in and out of the blood and lymphatic systems to defend against pathogens. Similarly, cancer cells move in and out of the blood and lymphatic systems before seeding metastatic sites. Both cell types are more invasive if they undergo a phenotypic transition from mesenchymal to amoeboid migration in response to mechanochemical cues [1]. Confinement has been shown to potently trigger the phenotypic transition from mesenchymal to amoeboid migration; however, the molecular mechanism(s) that underlie this phenomenon are not fully understood [2–4].

In melanoma and other cancers, confinement can trigger a phenotypic transition from mesenchymal to amoeboid migration. During mesenchymal migration, actin polymerization drives protrusion of the leading edge, whereas during amoeboid migration, protrusion of the leading edge is driven by blebs [5]. Blebs form when a segment of the plasma membrane separates from the underlying cortical actomyosin cytoskeleton, which requires high levels of intracellular pressure [6]. Consequently, cells with high levels of cortical actomyosin contractility are prone to blebbing [2]. In most cells, blebs are retracted within a minute [7]. Under certain conditions, however, cortical actomyosin flow in stable blebs drives “fast amoeboid migration” [2].

In vertical or 2-dimensional (2D) confinement, cell movement may be driven by cortical actomyosin flow in stable blebs [2–4]. As cells move in the direction of this bleb, it was termed a ‘leader bleb’ [4]. Accordingly, fast amoeboid migration has also been described as leader bleb-based migration (LBBM) [4]. Importantly, melanoma and other cancer cell types have been observed *in vivo* to use a phenotypically similar mode of migration by intravital microscopy [8]. Previous work by others demonstrated that confining cells down to 3 µm and low adhesion potently triggers the phenotypic transition to fast amoeboid (leader bleb-based) migration in transformed cells [2, 9]. Thus, elucidating the molecular mechanism(s) that underlie this phenomenon may provide new therapeutic avenues for the prevention of metastasis.

To undergo a phenotypic transition to amoeboid migration, cells must sense mechanochemical cues from the microenvironment. Work by others has shown that Piezo1, Phosphodiesterase 1 (PDE1), and cytosolic phospholipase A2 (cPLA2) promote migration in confined environments [10–12]. The plasma membrane tension sensor, Piezo1, was shown to promote migration through the Ca^2+^ mediated activation of PDE1 [10]. cPLA2 liberates arachidonic acid from the nuclear membrane, which activates myosin ATPases [11, 12]. For full activity, cPLA2 requires that the nuclear membrane be stretched [13, 14]. In this model, therefore, nuclear membrane stretch is key to promote migration [11, 12]. It remains unclear, however, if these factors are sufficient to induce a phenotypic switch from mesenchymal to amoeboid migration in confined environments.

Here, we demonstrate that melanoma cells sense increasing levels of confinement using the stretch activated cation selective channel, Piezo1. Upon inhibiting Piezo1 or Ca^2+^ chelation, melanoma cells less frequently adopt amoeboid migration in 2D and 3D (*i.e.*, microchannel) confined environments. In undulating channels, which are more likely to resemble the tortuous vasculature surrounding tumors, nearly all melanoma cells adopted an amoeboid mode of migration in a Piezo1/Ca^2+^ dependent fashion. Importantly, Piezo1/Ca^2+^ is found to activate inverted formin-2 (INF2), which induces actin cytoskeletal remodeling and de-adhesion. By promoting de-adhesion, the activation of INF2 by Piezo1/Ca^2+^ drives mesenchymal to amoeboid transition (MAT) in confined environments.

## Results

To promote fast amoeboid (leader bleb-based) migration, invasive melanoma (A375-M2) cells were placed in 2D confinement. To accomplish this, cells are placed under a slab of polydimethylsiloxane (PDMS), which is held at a defined height above cover glass by micron-sized beads [15]. Previously, it was shown that the switch to leader bleb-based migration (LBBM) occurs most frequently when cells are confined down to 3 µm [2]. Therefore, the PDMS was held above cover glass by 3 µm beads [15]. To block integrin adhesion, the PDMS and cover glass were coated with Bovine Serum Albumin (BSA; 1%). Under these conditions, melanoma cells migrate in the direction of a leader bleb (Figure 1A, Figure 1-video 1). These cells are termed leader mobile (LM). Cells that form leader blebs but do not move are referred to as leader non-mobile (LNM; Figure 1-figure supplement 1A, Figure 1-video 2). Cells without a leader bleb and do not move are termed no leader (NL; Figure 1-figure supplement 1A, Figure 1-video 3). To evaluate the role of Ca^2+^ in regulating LBBM, we chelated Ca^2+^ using BAPTA (10 µM). This treatment resulted in an over 2-fold reduction in the number of leader mobile cells (Figure 1B). Therefore, we wondered if Piezo1 and/or 2 are required in melanoma cells for LBBM. Using RT-qPCR, we could only detect Piezo1, therefore we focused on Piezo1 for the rest of the study (Figure 1C, Figure 1-figure supplement 2). To determine if Piezo1 has a role in sensing confinement, we correlated the level of intracellular Ca^2+^ with height. In melanoma cells treated with a non-targeting, *i.e.*, control, small interfering RNA (siRNA), intracellular Ca^2+^ was found to sharply increase with confinement (Figure 1D). In contrast, intracellular Ca^2+^ levels and confinement were poorly correlated in cells lacking Piezo1 (Figure 1E-F). Upon evaluating migration in 2D confinement, we observed an over 2-fold reduction in the number of leader mobile cells (Figure 1G-H). Similarly, using a Piezo1 inhibitor, the percent and speed of leader mobile cells were significantly reduced (Figure 1-figure supplement 3A-B). Whereas the percentage and speed of leader mobile cells was significantly increased by a Piezo1 activator (Figure 1-figure supplement 3C-D). Using our Fiji (https://fiji.sc/) plugin, Analyze_Blebs, we found that the largest bleb area, aspect ratio, and roundness were reduced and increased by a Piezo1 inhibitor and activator, respectively (Figure 1-figure supplement 4A-F) [16]. Thus, Piezo1/Ca^2+^ promotes LBBM in 2D confinement.

**Figure 1.**
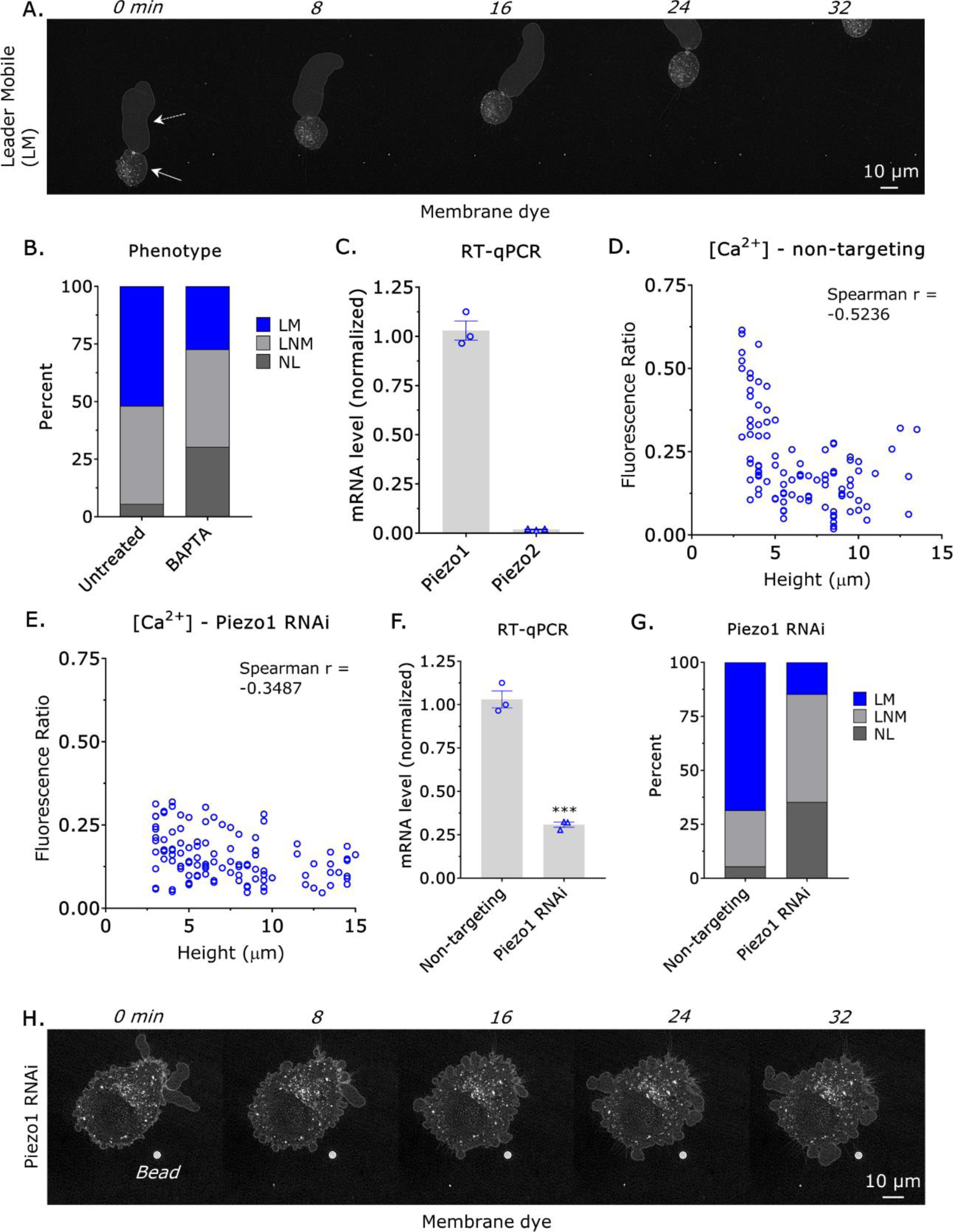
Piezo1/Ca^2+^ promotes fast amoeboid (leader bleb-based) migration. **A.** Montage of a leader mobile (LM) cell vertically confined down to a height of 3 µm. Cells were stained with a far-red membrane dye. Arrows point to the cell body (solid) and leader bleb (dashed), which is a large and stable bleb. **B.** Percent of untreated (n=54) or BAPTA (10 µM; n=33) treated cells adopting a leader mobile (LM), leader non-mobile (LNM), or no leader (NL) phenotype (χ^2^ ≤ 0.0001). **C.** RT-qPCR for Piezo1 and 2 on total mRNA (mean +/- SEM). **D.-E.** Compared to cells treated with a non-targeting siRNA, intracellular Ca^2+^ levels in cells treated with a Piezo1 siRNA no longer correlates with confinement height. Intracellular Ca^2+^ levels were measured by taking the fluorescence ratio of a far-red fluorescent Ca^2+^ indicator to a green fluorescent dye. **F.** 3 days after transfection, Piezo1 RNAi was confirmed by RT-qPCR on total mRNA (Student’s t-test; mean +/- SEM). **G.** Percent of non-targeting (n=73) or Piezo1 siRNA (n=34) treated cells adopting a leader mobile (LM), leader non-mobile (LNM), or no leader (NL) phenotype (χ^2^ ≤ 0.0001). **H.** Montage of a no leader (NL) cell vertically confined down to 3 µm treated with a Piezo1 siRNA. Cells were stained with a far-red membrane dye. PDMS was coated with Bovine Serum Albumin (BSA; 1%). * - p ≤ 0.05, ** - p ≤ 0.01, *** - p ≤ 0.001, and **** - p ≤ 0.0001

Within tissues, cells may encounter channel-like environments, such as within micro-capillary/lymphatic vessels [1]. Therefore, we then placed invasive melanoma (A375-M2) cells in microfabricated channels coated with VCAM-1 (1 µg/mL). Under these conditions, cells predominantly adopt a hybrid mode of migration, which has hallmarks of mesenchymal and amoeboid migration (*e.g.,* blebs) (Figure 2A, Figure 2-video 1). Notably, we have also observed melanoma cells adopting a hybrid mode of migration in fibronectin coated (10 µg/mL) channels [17]. In the presence of a Piezo1 inhibitor, however, cells predominantly adopted a mesenchymal mode of migration (Figure 2B). We then wondered if we could bias cells towards an amoeboid phenotype using undulating channels, which are more likely to resemble the tortuous vasculature surrounding tumors. We speculate that in undulating channels, Piezo1 may be gated open through a curvature induced increase in plasma membrane tension. Indeed, in undulating channels, we found that cells were almost entirely amoeboid (Figure 2C, Figure 2-video 2). If Ca^2+^ was chelated with BAPTA (10 µM), however, cells predominantly adopted a mesenchymal mode of migration (Figure 2D). Notably, mesenchymal cells were also frequently motile (Figure 2D). In agreement with these data, intracellular Ca^2+^ levels were elevated in cells within undulating channels (Figure 2E). Treating cells with a Piezo1 inhibitor led to some cells adopting a hybrid phenotype (Figure 3A-B). Piezo1 RNAi, however, led to a significant increase in the number of cells adopting a mesenchymal mode of migration (Figure 3C). Interestingly, Piezo1 RNAi had no effect on the speed and directionality of motile cells; therefore, Piezo/Ca^2+^ appears to primarily promote the appearance of amoeboid features (*e.g.*, blebs) in channels (Figure 3D-E).

**Figure 2.**
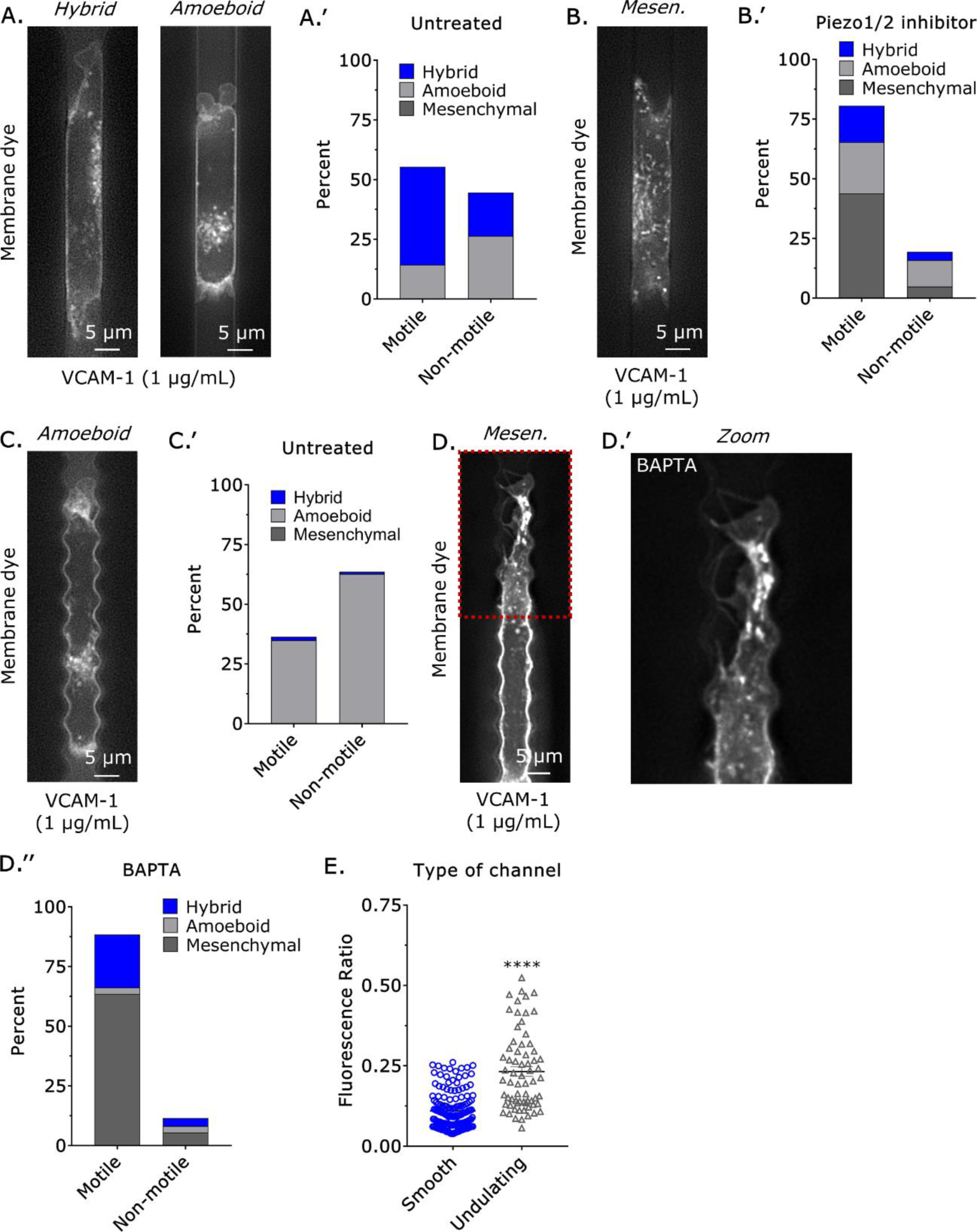
In microchannels, motile cells predominantly adopt a hybrid phenotype. **A.** In microchannels, cells predominantly adopt a hybrid phenotype (n=28), which has hallmarks of amoeboid (*i.e.*, blebs) and mesenchymal migration. **B.** In the presence of a Piezo1/2 inhibitor (GsMTx4; 10 µM) (n=165), cells predominantly adopt a mesenchymal phenotype (χ^2^ ≤ 0.0001). **C.** In undulating channels, which are more likely to resemble the tortuous vasculature surrounding tumors, cells predominantly adopt an amoeboid phenotype (n=144). **D.** In undulating channels, cells treated with the Ca^2+^ chelator, BAPTA (10 µM; n=148), predominantly adopt a mesenchymal phenotype (χ^2^ ≤ 0.0001). No blebs are observed in the zoomed image (D’). **E.** Compared to cells in smooth channels, intracellular Ca^2+^ levels are elevated in cells within undulating channels. Intracellular Ca^2+^ levels were measured by taking the fluorescence ratio of a far-red fluorescent Ca^2+^ indicator to a green fluorescent dye (Student’s t-test; mean +/- SEM). Microchannels are coated with VCAM-1 (1 µg/mL) and are 8 µm in height, 8 µm in width, and 100 µm in length.

**Figure 3.**
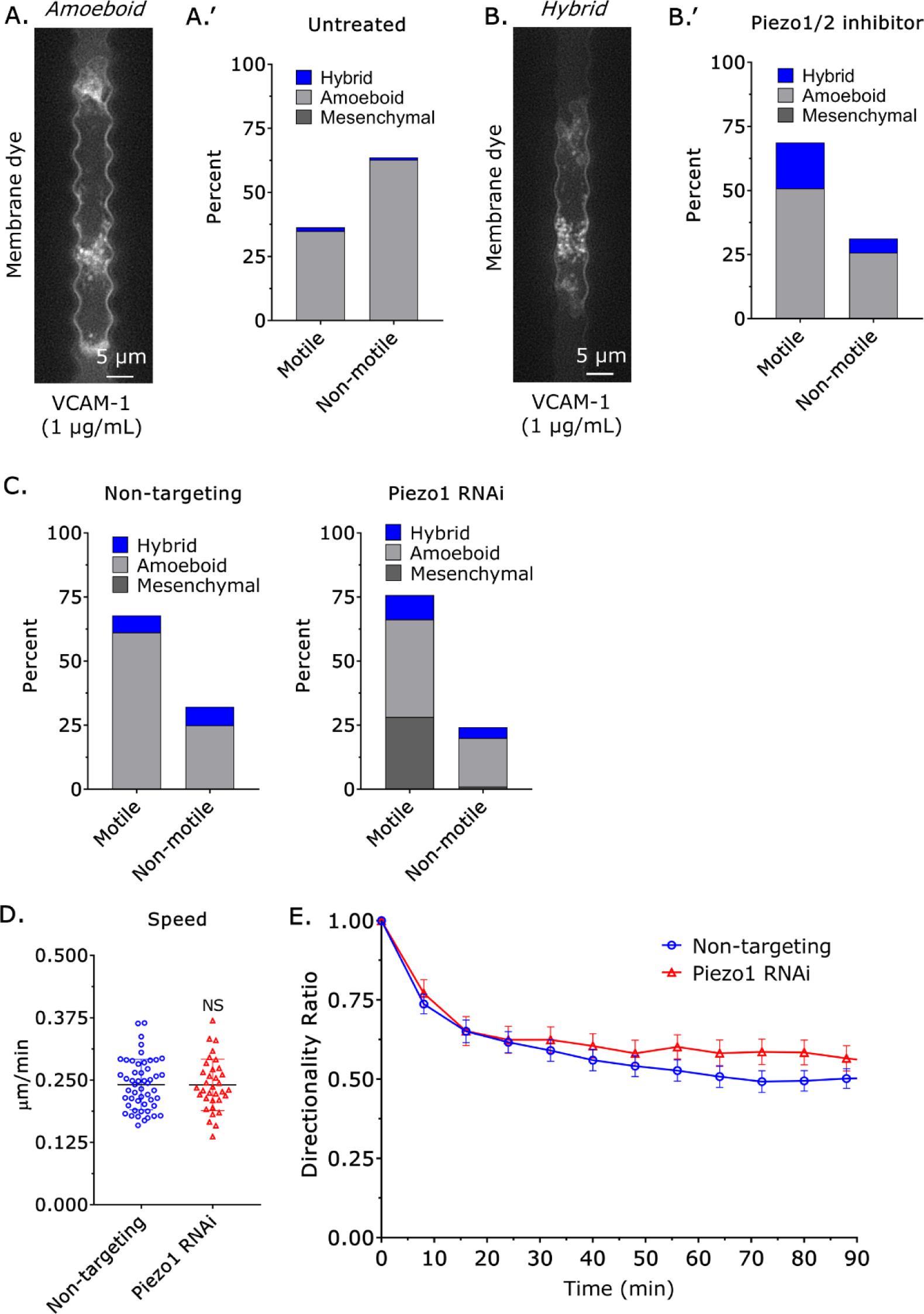
Piezo1 promotes amoeboid migration in undulating channels. **A.-B.** In undulating channels, cells treated with a Piezo1 inhibitor (GsMTx4; 10 µM) (n=96) more often adopt a hybrid phenotype (χ^2^ ≤ 0.0001). For comparison, figure 2C is re-displayed. **C.** Compared to cells treated with a non-targeting siRNA (n=155), cells treated with a Piezo1 siRNA (n=210) more often adopt a mesenchymal phenotype (χ^2^ ≤ 0.05). **D.** Based on manual cell tracking, the speed of non-targeting and Piezo1 siRNA treated cells is similar (Student’s t-test; mean +/- SEM). **E.** Based on manual cell tracking, the directionality ratio for non-targeting (n=51) and Piezo1 siRNA (n=35) treated cells is similar (mean +/- SEM). Microchannels are coated with VCAM-1 (1 µg/mL) and are 8 µm in height, 8 µm in width, and 100 µm in length.

Intracellular Ca^2+^ regulates several signaling pathways, including cytosolic phospholipase A2 (cPLA2). For full activity, however, cPLA2 requires nuclear membrane tension [13, 14]. Accordingly, through the liberation of arachidonic acid from nuclear membranes, cPLA2 has been shown to promote migration in confined environments [11, 12]. Therefore, we wondered if cPLA2 may be downstream of Piezo1/Ca^2+^. In undulating channels, we could not detect any difference in the number or phenotype of motile cells after cPLA2 RNAi (PLA2G4A; Figure 4A). Moreover, cPLA2 RNAi had no significant effect on the speed and directionality of motile cells (Figure 4-figure supplement 1A-B). We then set out to determine if either myosin light chain kinase (MYLK), which is activated by Ca^2+^/calmodulin, or Rho-associated protein kinase (ROCK) may instead regulate amoeboid migration downstream of Piezo1/Ca^2+^. By RT-qPCR, we could detect the MYLK, ROCK1, and ROCK2 isoforms in cells (Figure 4B). Moreover, each isoform could be depleted by RNAi (Figure 4C). Surprisingly, in undulating channels, only RNAi of ROCK2 led to a significant change in the phenotype of motile cells. More specifically, cells predominantly adopted a mesenchymal mode of migration after ROCK2 RNAi, whereas non-targeting, *i.e.*, control, MYLK, and ROCK1 RNAi cells predominantly adopted an amoeboid mode of migration (Figure 4D). Notably, MYLK, ROCK1, and ROCK2 RNAi had no significant effect on the speed or directionality of motile cells (Figure 5-figure supplement 1A-B). As ROCK2 has been reported to have functions in the cytosol and nucleus (*i.e.*, regulating gene transcription), we determined if upregulating myosin ATPase activity is sufficient to restore an amoeboid phenotype after RNAi. By dephosphorylating the regulatory light chain (RLC), myosin phosphatase 1 (MYPT1) decreases myosin ATPase activity in cells [18]. We determined, therefore, if MYPT1 RNAi is sufficient to restore an amoeboid phenotype. In contrast to cells with a ROCK2 siRNA alone, cells with a ROCK2 and MYPT1 siRNA (*i.e.*, ROCK2 + MYPT1 RNAi) predominantly adopted an amoeboid phenotype (Figure 5A-E). In 2D confinement, we observed similar effects. More specifically, we observed fewer leader mobile (LM) cells after treatment with a ROCK2 siRNA, whereas treatment with a ROCK2 and MYPT1 siRNA had the opposite effect (Figure 5-figure supplement 2A). Moreover, cPLA2 RNAi had no effect on the number of leader mobile (LM) cells (Figure 5-figure supplement 2B). Using our Fiji (https://fiji.sc/) plugin, Analyze_Blebs, we could detect changes in shape descriptors after ROCK2 RNAi, including a decrease in the largest bleb area (Figure 5-figure supplement 3A-F) [16]. Thus, ROCK2 biases cells towards an amoeboid phenotype in confined environments.

**Figure 4.**
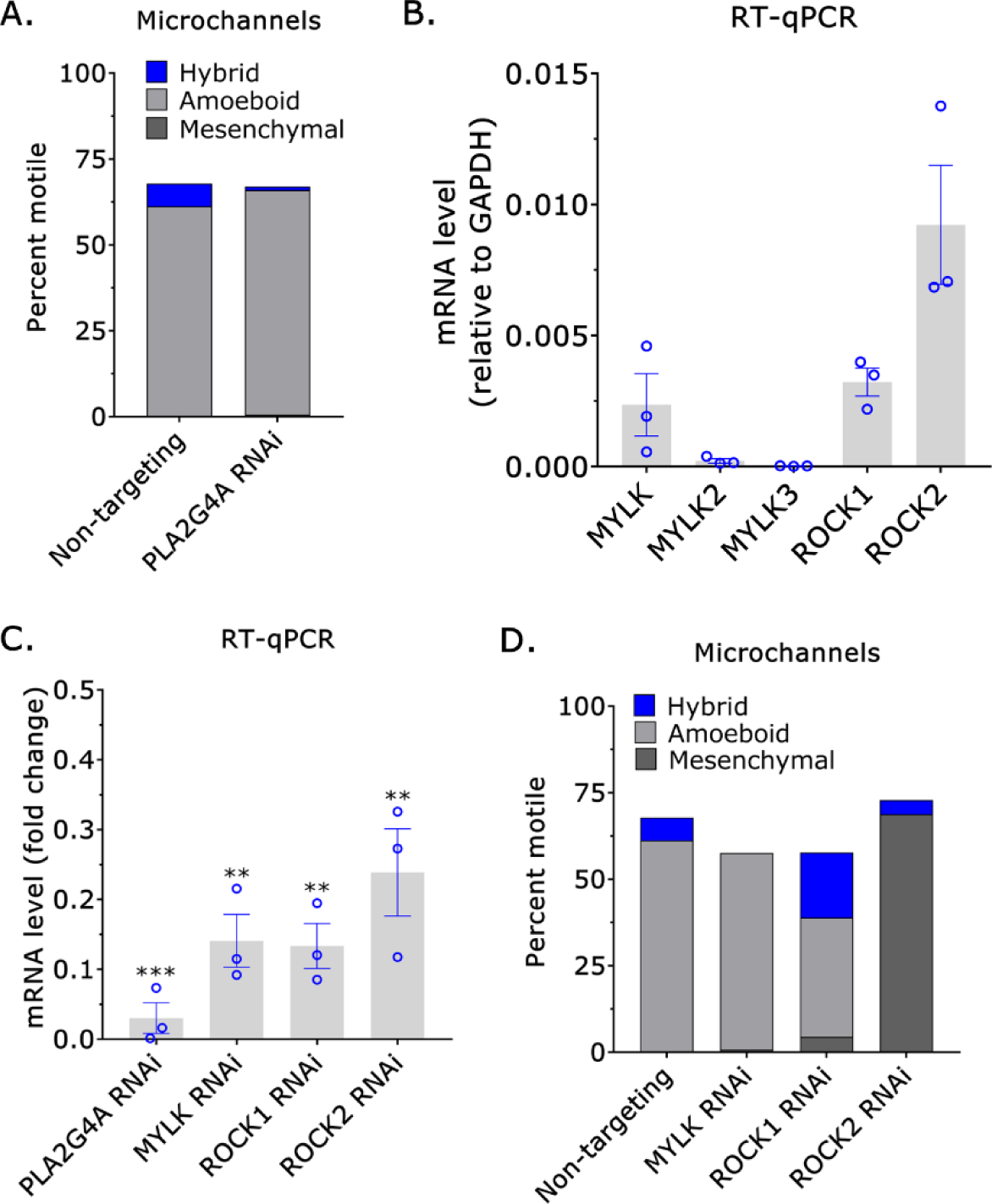
ROCK2 promotes amoeboid migration in undulating channels. **A.** Compared to non-targeting (n=155), the percent of motile cells or their phenotype is not significantly affected by treatment with a cPLA2 siRNA (PLA2G4A; n=135) (χ^2^=NS). **B.** RT-qPCR for MYLK, MYLK2, MYLK3, ROCK1, and ROCK2 on total mRNA (mean +/- SEM). **C.** 3 days after transfection, RNAi of cPLA2 (PLA2G4A), MYLK, ROCK1, and ROCK2 was confirmed by RT-qPCR on total mRNA (Student’s t-test; mean +/- SEM). **D.** Compared to non-targeting (n=135), MYLK (n=146; χ^2^=NS), and ROCK1 (n=161; χ^2^ ≤ 0.0001), ROCK2 siRNA (n=170; χ^2^ ≤ 0.0001) treated cells were predominantly mesenchymal, whereas the percent of motile cells was not significantly affected by any siRNA. Microchannels are coated with VCAM-1 (1 µg/mL) and are 8 µm in height, 8 µm in width, and 100 µm in length. * - p ≤ 0.05, ** - p ≤ 0.01, *** - p ≤ 0.001, and **** - p ≤ 0.0001

**Figure 5.**
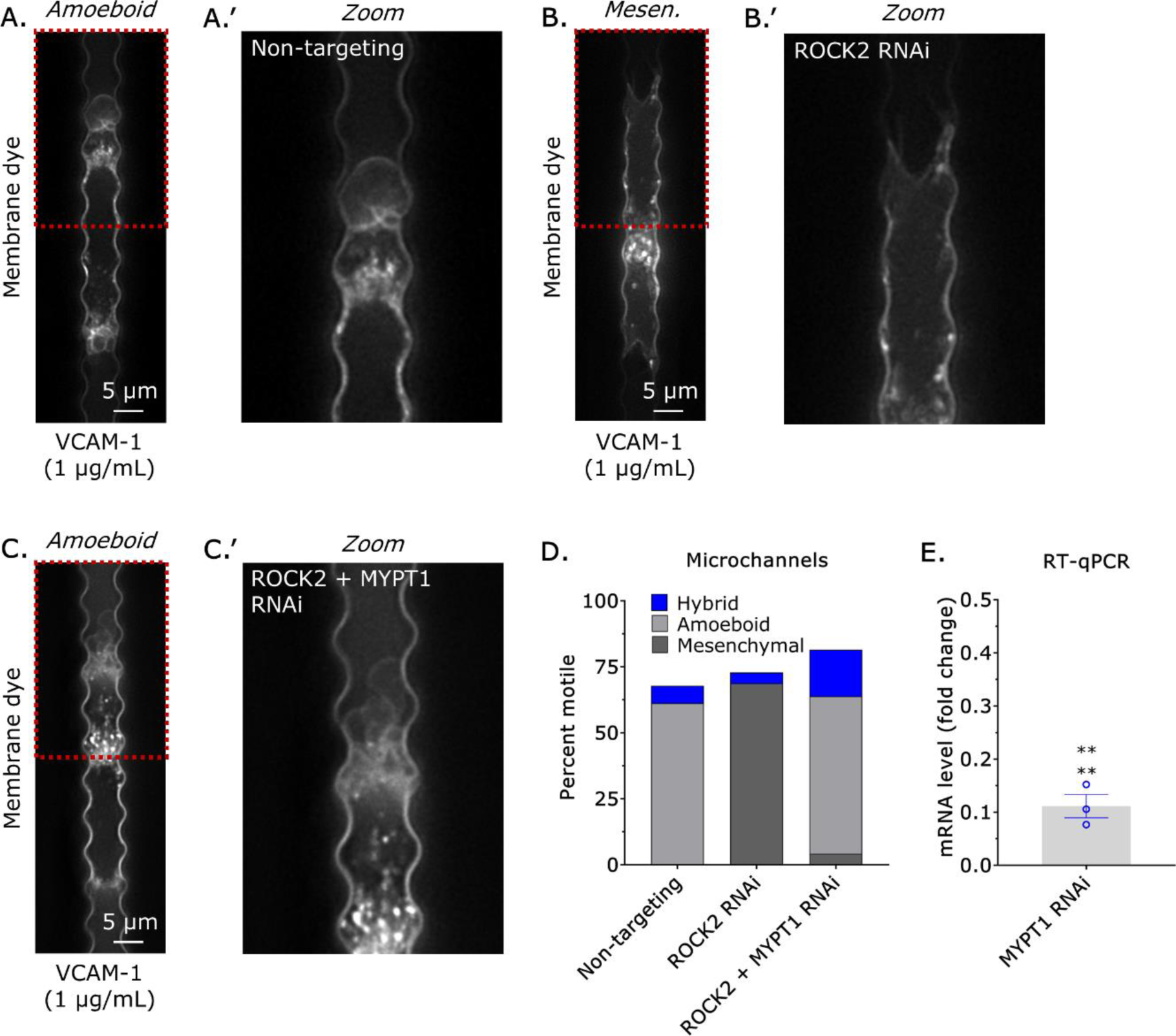
ROCK2 promotes amoeboid migration in undulating channels through increasing actomyosin contractility. **A.** Cells treated with a non-targeting siRNA (n=155) predominantly adopt an amoeboid phenotype. Blebs are observed in the zoomed image (A’). **B.** Cells treated with a ROCK2 siRNA (n=170) predominantly adopt a mesenchymal phenotype. No blebs are observed in the zoomed image (B’). **C.** Compared to cells treated with a ROCK2 siRNA (n=170) alone, cells treated with both a ROCK2 and MYPT1 siRNA (n=223) predominantly adopt an amoeboid phenotype. Blebs are observed in the zoomed image (C’). **D.** Percent motile and phenotype of cells treated with a non-targeting (n=94), ROCK2 (n=123; χ^2^ ≤ 0.0001), and with both a ROCK2 and MYPT1 siRNA (n=182; χ^2^=NS). **E.** 3 days after transfection, RNAi of MYPT1 was confirmed by RT-qPCR on total mRNA (Student’s t-test; mean +/- SEM). Microchannels are coated with VCAM-1 (1 µg/mL) and are 8 µm in height, 8 µm in width, and 100 µm in length. * - p ≤ 0.05, ** - p ≤ 0.01, *** - p ≤ 0.001, and **** - p ≤ 0.0001

So far, we have demonstrated that Piezo1/Ca^2+^ and ROCK2 bias cells towards an amoeboid phenotype in confined environments. We still wondered, however, what factors may be directly downstream of Piezo1/Ca^2+^. Using a ROCK2 biosensor (Eevee-ROCK), we could not detect any increase in ROCK2 kinase activity upon activating Piezo1 (Figure 6-figure supplement 1A-C) [19]. Thus, the prototypical regulators of myosin ATPase activity may not be regulated by Piezo1/Ca^2+^ in this cell type. We determined, therefore, if there are actin cytoskeletal remodeling factors downstream of Piezo1/Ca^2+^ that could bias cells towards an amoeboid phenotype. A phenomenon termed, calcium-mediated actin reset or CaAR, occurs following activation of inverted formin-2 (INF2) [20]. During CaAR, actin is remodeled or “reset” by INF2, which is an atypical formin activated by Ca^2+^ that can be endoplasmic reticulum (ER) localized [20, 21]. Possibly due to monomer sequestration, the activation of INF2 by Ca^2+^ leads to a rapid de- and then re-polymerization of actin (*i.e.*, reset) throughout the cell [20]. Using RT-qPCR, we could detect several formins, including INF2 in invasive melanoma (A375-M2) cells (Figure 6A). As measured by LifeAct-mEmerald, we could measure a rapid increase in F-actin near the nucleus after treatment with a Piezo1 activator (Figure 6B-C, Figure 6-video 1) [22]. Simultaneously, we could detect a decrease in F-actin elsewhere in cells, such as near the cell edge (Figure 6B-C, Figure 6-video 1). After INF2 RNAi, however, the increase and decrease in F-actin near the nucleus and cell edge, respectively, after treatment with a Piezo1 activator were blunted (Figure 6B-C, Figure 6-video 2). Interestingly, the peak [Ca^2+^] elicited by a Piezo1 activator was reduced by ∼25% in cells treated with an INF2 siRNA (Figure 6-figure supplement 2A). However, Ca^2+^ levels in non-targeting and INF2 siRNA treated cells were equivalent by ∼5 min (Figure 6-figure supplement 2A). Collectively, these data establish that INF2 remodels the actin cytoskeleton in response to Piezo1/Ca^2+^.

**Figure 6.**
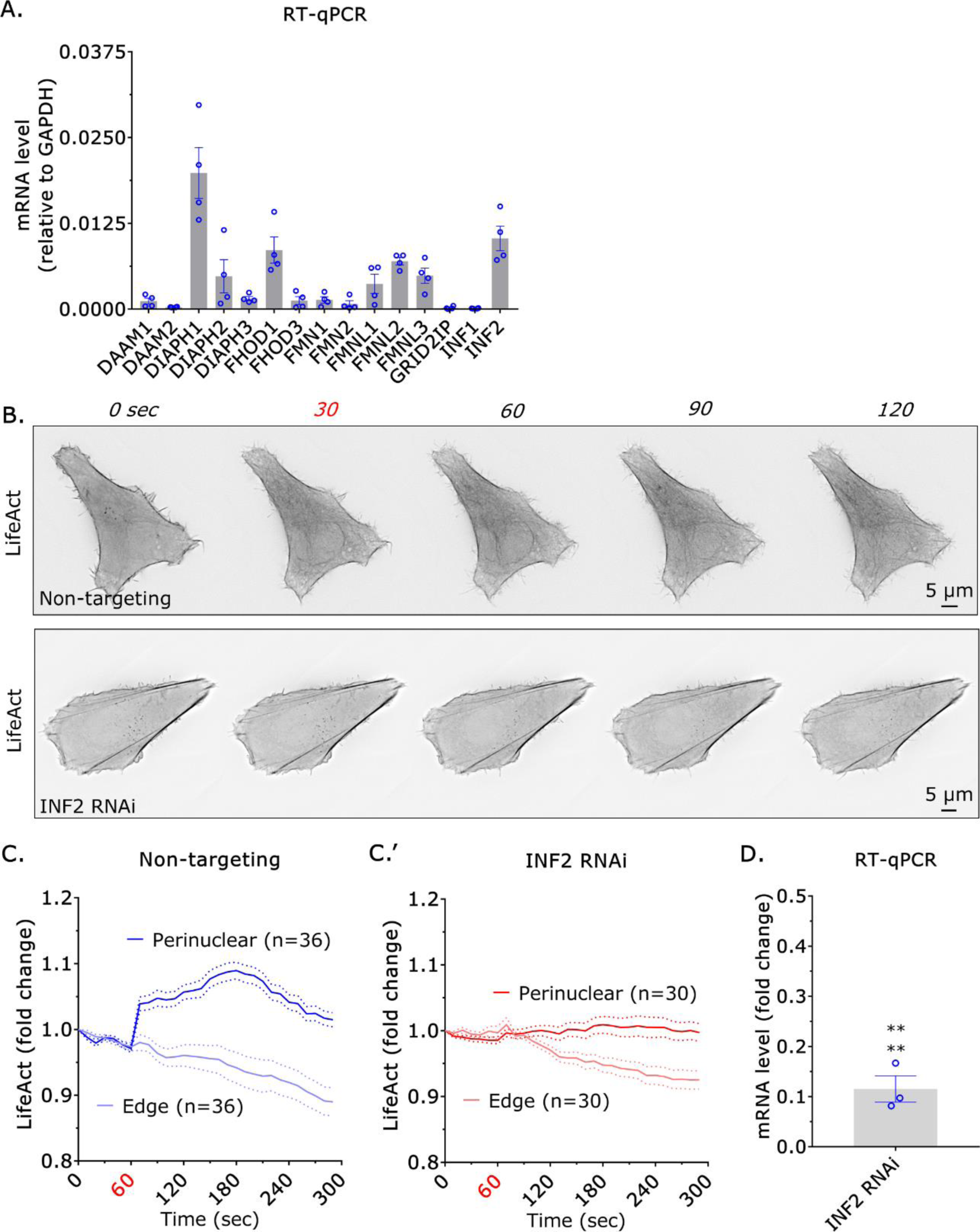
INF2 remodels the actin cytoskeleton downstream of Piezo1/Ca^2+^. **A.** RT-qPCR for mammalian formins on total mRNA (mean +/- SEM). **B.** Montages of cells transiently transfected with a non-targeting or INF2 siRNA and LifeAct-mEmerald treated with a Piezo1/2 activator (Yoda1; 20 µM) at 30 sec. **C.** Plot of LifeAct-mEmerald fluorescence intensities over time in cells treated with a Piezo1/2 activator (Yoda1; 20 µM) at 60 sec (mean +/- SEM). Fluorescence intensities were measured within regions of interest near the nucleus or cell edge. Cells were treated with a non-targeting (C) or INF2 (C’) siRNA. **D.** 3 days after transfection, RNAi of INF2 was confirmed by RT-qPCR on total mRNA (Student’s t-test; mean +/- SEM). Cells were plated on fibronectin coated (10 µg/mL) glass. * - p ≤ 0.05, ** - p ≤ 0.01, *** - p ≤ 0.001, and **** - p ≤ 0.0001

Having established that INF2 remodels the actin cytoskeleton in response to Piezo1/Ca^2+^, we wondered if INF2 could bias cells towards an amoeboid phenotype in confined environments. In undulating channels, invasive melanoma (A375-M2) cells treated with a non-targeting, *i.e.*, control, siRNA predominantly adopted an amoeboid phenotype, whereas cells treated with an INF2 siRNA predominantly adopted a mesenchymal phenotype (Figure 7A-B, Figure 7-video 1). After INF2 RNAi, cells were also faster and more directional (Figure 7C-D). In the absence of VCAM-1 (1 µg/mL), however, cells even without INF2 predominantly adopt an amoeboid migration mode (Figure 7E). In 2D confinement, we also did not observe any difference in the number of cells adopting a leader mobile (LM) phenotype after INF2 RNAi (Figure 7-figure supplement 1A). As INF2 RNAi has no effect in non-adherent environments, INF2 may drive amoeboid migration by promoting de-adhesion. Indeed, in cells treated with a non-targeting siRNA, we could observe a decrease in Vinculin, which marks focal adhesions, after treatment with a Piezo1 activator (Figure 7F). After INF2 RNAi, however, Vinculin levels were unchanged after treatment with a Piezo1 activator (Figure 7F). These results were confirmed by cell impedance measurements, which increases with cell adhesion. More specifically, cells treated with a non-targeting, *i.e.*, control, siRNA de-adhered shortly after treatment with a Piezo1 activator, whereas cells treated with an INF2 siRNA failed to de-adhere (Figure 8A). Freshly plated cells also displayed superior adhesion after INF2 RNAi, as measured by cell impedance (Figure 8B). By inducing actin cytoskeletal remodeling and de-adhesion, INF2 may drive mesenchymal to amoeboid transition (MAT) in confined environments. To test this idea, we combined 2D confinement with adhesive (fibronectin; 10 µg/mL) micropatterns. When confined down to 3 µm, cells treated with a non-targeting, *i.e.*, control, siRNA frequently crossed adhesive patterns using fast amoeboid (leader bleb-based) migration (Figure 8C, Figure 8-video 1).

**Figure 7.**
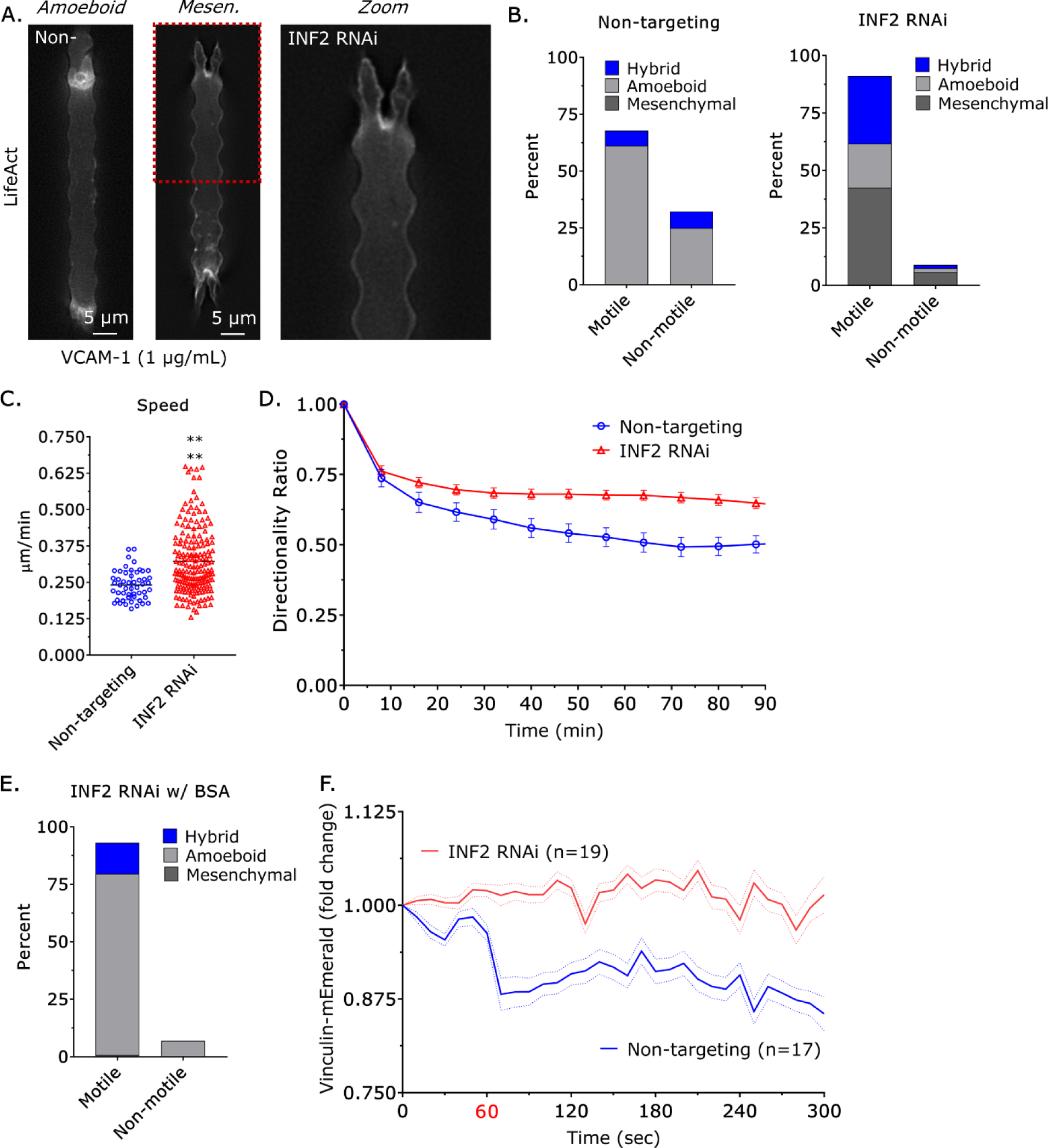
INF2 promotes amoeboid migration in undulating channels downstream of Piezo1/Ca^2+^. **A.** Representative images of cells transiently transfected with non-targeting or an INF2 siRNA and LifeAct-mEmerald. No blebs can be observed in the zoomed image (INF2 RNAi). **B.** Compared to non-targeting (n=155), INF2 siRNA (n=250) treated cells were predominantly mesenchymal (χ^2^ ≤ 0.0001). **C.** Compared to non-targeting, INF2 siRNA treated cells are significantly faster (Student’s t-test; mean +/- SEM). **D.** Compared to non-targeting (n=51), INF2 siRNA (n=192) treated cells migrate more directionally (mean +/- SEM). **E.** In undulating channels coated with BSA (1%), which blocks adhesion, cells treated with an INF2 siRNA (n=107) predominantly adopt an amoeboid phenotype. **F.** Compared to non-targeting, INF2 siRNA treated cells plated on VCAM-1 (1 µg/mL) coated glass do not display a decrease in Vinculin-mEmerald after treatment with a Piezo1/2 activator (Yoda1; 20 µM) at 60 sec (mean +/- SEM). Vinculin-mEmerald was plotted as a ratio of fluorescence at focal adhesions to an uninvolved region. Microchannels are coated with VCAM-1 (1 µg/mL) and are 8 µm in height, 8 µm in width, and 100 µm in length. * - p ≤ 0.05, ** - p ≤ 0.01, *** - p ≤ 0.001, and **** - p ≤ 0.0001

**Figure 8.**
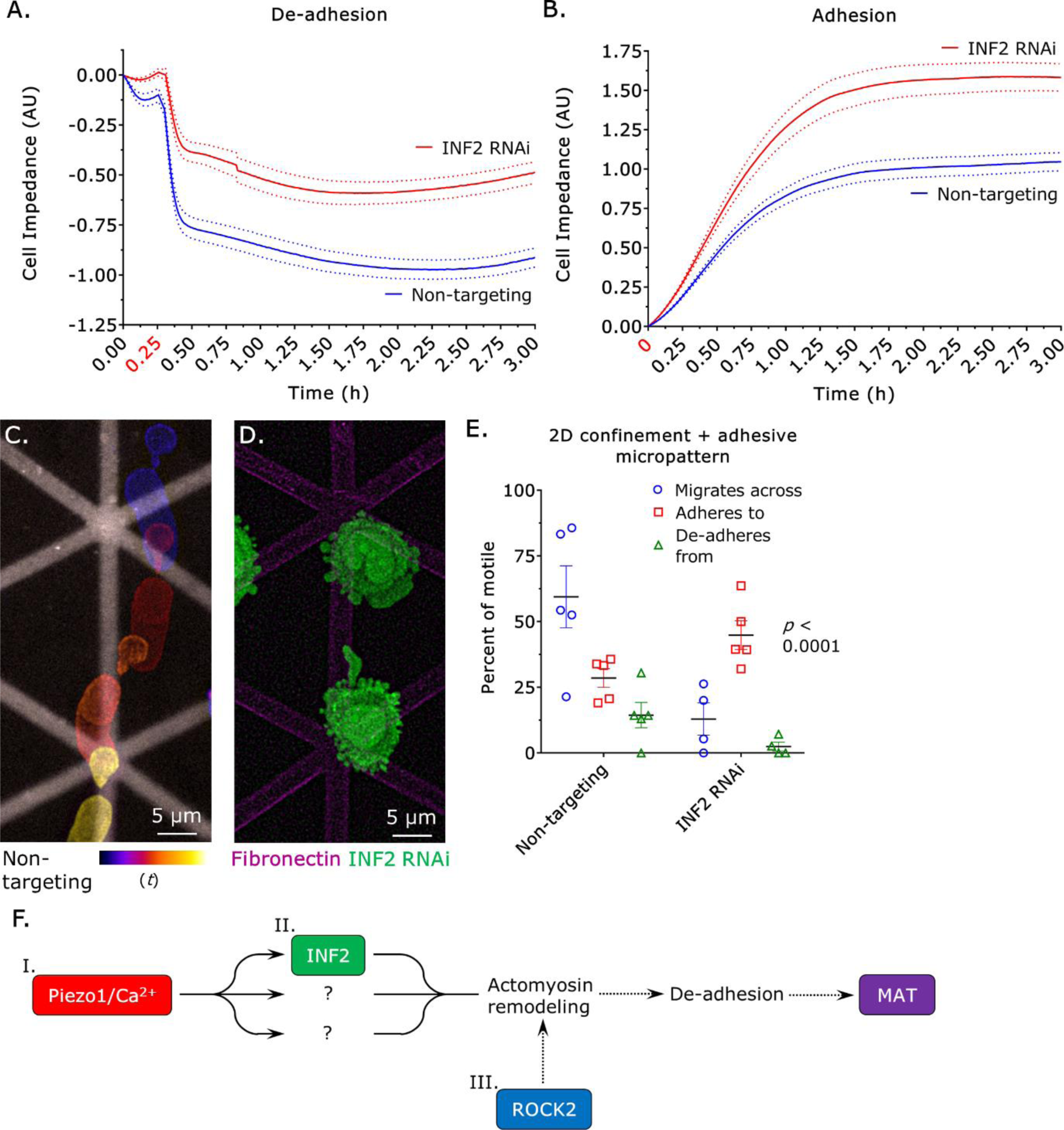
INF2 drives amoeboid migration by promoting de-adhesion. **A.** Compared to non-targeting, INF2 siRNA treated cells fail to de-adhere from VCAM-1 (1 µg/mL) coated cover glass after treatment with a Piezo1/2 activator (Yoda1; 20 µM) at 15 min (mean +/- SEM). Cell impedance was measured using a Real Time Cell Analyzer (RTCA; Agilent). **B.** Compared to non-targeting, INF2 siRNA treated cells adhere faster to VCAM-1 (1 µg/mL) coated cover glass (mean +/- SEM). Cell impedance was measured using a Real Time Cell Analyzer (RTCA; Agilent). **C.** Temporal color-code image of a non-targeting siRNA treated cell migrating across an adhesive (fibronectin; 10 µg/mL) micropattern. Cells were confined down to 3 µm using a dynamic cell confiner (4Dcell). **D.** INF2 siRNA (green) treated cells adhere to adhesive (fibronectin; 10 µg/mL) micropatterns (magenta). Cells were confined down to 3 µm using a dynamic cell confiner (4Dcell). **E.** Percent of cells that migrate across patterns (circles), randomly migrate into and then adheres to the pattern (squares), and that de-adheres from the pattern and then migrates away (triangles) for non-targeting (n=247) and INF2 (n=249) siRNA treated cells. A p-value comparing non-targeting to INF2 siRNA treated cells was calculated using a one-way ANOVA. **F.** In the proposed model, Piezo1 is opened by plasma membrane tension (I), intracellular Ca^2+^ activates INF2 and possibly other factors (II), and ROCK2 phosphorylates the regulatory light chain of myosin, myosin phosphatase, LIMK1/2, and others (III) to induce actomyosin remodeling, de-adhesion, and mesenchymal to amoeboid transition (MAT).

Despite being confined down to 3 µm, cells were frequently found adhered to the pattern after INF2 RNAi (Figure 8D, Figure 8-video 2). Consistent with this observation, we more frequently observed cells treated with an INF2 siRNA randomly migrating into and then adhering to the pattern (Figure 8E). Additionally, INF2 siRNA treated cells were less likely to de-adhere from the pattern and then migrate away (Figure 8E). Collectively, our data demonstrates that cells require the activation of INF2 by Piezo1/Ca^2+^ for phenotypic plasticity. Thus, enabling the movement of cells through environments with challenging mechanochemical properties.

## Discussion

Disseminating tumor cells must adapt to diverse mechanochemical environments before seeding metastatic sites. Similarly, immune cells must adapt to diverse mechanochemical environments to defend against pathogens. Both immune and cancer cells have been shown to adopt amoeboid modes of migration under conditions of low adhesion and confinement [2, 23]. Among cancer cell types, melanoma represents the best studied in terms of amoeboid migration [24]. Notably, human melanoma cells have been observed migrating in the direction of a large bleb within tumors implanted in immunocompromised mice by intravital imaging [8].

Moreover, in sectioned tumors from melanoma patients, predominantly amoeboid cells are observed at invasive fronts [25]. Accordingly, the dissemination of melanoma tumors is likely promoted by amoeboid migrating cells. Although candidate factors have been identified, it remains unclear what drives amoeboid migration [10–12]. In a Piezo1 dependent fashion, we show that intracellular [Ca^2+^] increases as cells are vertically confined. Similarly, inhibiting or activating Piezo1 led to a decrease and increase in leader bleb-based migration (LBBM), respectively. Consistent with its effects on LBBM, Piezo1 activity also correlated with bleb size. In zebrafish embryos, cytosolic phospholipase A2 (cPLA2) was shown to promote a phenotypically similar mode of migration [12]. Here, we were unable to detect any difference in the rate of LBBM after cPLA2 RNAi (Figure 5-figure supplement 2B). In transmigration assays, however, we previously reported a defect in migration through confining pores [26]. Thus, cPLA2 appears to play a nuanced role during confined migration. In contrast, ROCK2 was identified as being essential for LBBM, which is consistent with work by others on amoeboid migrating cells at the invasive fronts of melanoma tumors [27].

In microchannels coated with VCAM-1, cells predominantly adopted a hybrid phenotype. In contrast to cells confined in 2D, which form leader blebs, cells in VCAM-1 coated microchannels form F-actin and intracellular pressure (*i.e.*, blebs) driven protrusions. Like what was found for cells confined in 2D, however, the appearance of blebs was promoted by Piezo1/Ca^2+^. Thus, in 2D and 3D (*i.e.*, microchannels) confined environments, Piezo1 is gated open by plasma membrane tension. In undulating channels, which are more likely to resemble the tortuous vasculature surrounding tumors, cells predominantly adopted an amoeboid phenotype. These data are consistent with a curvature induced increase in plasma membrane tension. In agreement with this concept, more cells adopted a mesenchymal phenotype after Piezo1 RNAi. In comparison to Piezo1 RNAi, however, the chelation of Ca^2+^ with BAPTA had a much more potent effect. Therefore, it is possible that other Ca^2+^ selective channels contribute to this phenomenon. Recently, a mechanosensitive transient receptor potential (TRP) channel was shown to promote melanoma metastasis [28].

In cells, Ca^2+^ regulates several signaling pathways. Myosin light chain kinase (MYLK), for instance, is activated by Ca^2+^/calmodulin [29]. To our surprise, RNAi of MYLK had no effect on amoeboid migration. In contrast, we identified ROCK2 as essential for amoeboid migration. Therefore, we wondered if Piezo1/Ca^2+^ could activate ROCK2. Using a biosensor for ROCK (Eevee-ROCK), we were unable to detect any increase in ROCK2 kinase activity upon activating Piezo1 [19]. Accordingly, we looked to actin cytoskeletal remodeling proteins as candidate factors downstream of Piezo1/Ca^2+^. This led to us evaluating the role of inverted formin-2 (INF2), which was shown to promote calcium-mediated actin reset or CaAR [20]. In undulating channels, cells predominantly adopted a mesenchymal phenotype after INF2 RNAi. In the absence of VCAM-1, however, cells still adopted an amoeboid phenotype after INF2 RNAi. As INF2 RNAi had no effect in non-adherent environments, INF2 may drive amoeboid migration by promoting de-adhesion. Indeed, we found that Piezo1/Ca^2+^ induced de-adhesion, whereas cells failed to de-adhere after INF2 RNAi. Moreover, actin cytoskeletal remodeling in response to Piezo1/Ca^2+^ was abrogated in cells after RNAi of INF2. It is noteworthy that INF2 RNAi cells, which adopt a mesenchymal phenotype, migrated faster and more directionally. By promoting de-adhesion, INF2 may contribute to cancer metastasis by promoting mesenchymal to amoeboid transition (MAT). In agreement with this concept, cells were found to require INF2 for migration across micropatterned surfaces. In heterogenous tissues, disseminating tumor cells are likely to encounter low and highly adhesive environments. Having identified INF2 here, it remains likely that a suite of Ca^2+^ responsive factors is likely to regulate MAT in cell type specific fashions (Figure 8F).

The capacity for switching between mesenchymal and amoeboid migration modes is likely to increase the metastatic potential of melanoma tumors. Based on RNA sequencing data from The Cancer Genome Atlas (https://www.cancer.gov/tcga), high levels of Piezo1 and ROCK2 are correlated with poor survival across melanoma patients (hazard ratio > 2; Figure 8-figure supplement 1A-D, Figure 8-figure supplement 1-source data 1). Here, we have identified INF2 as an important mediator of phenotypic plasticity downstream of Piezo1/Ca^2+^ in invasive melanoma cells. Future work will aim to determine the range of mechanochemical environments in which INF2 is required for rapid migration in diverse cell types.

## Materials and Methods

### Cell culture

A375-M2 (CRL-3223) were obtained from the American Type Culture Collection (ATCC; Manassas, VA). Cells were cultured in high-glucose DMEM with pyruvate (Thermo Fisher, Carlsbad, CA) or FluoroBrite (cat no. A1896701; Thermo Fisher) supplemented with fetal bovine serum (cat no. 12106C; Sigma Aldrich, St. Louis, MO), L-glutamine (cat no. 35050061; Thermo Fisher), antibiotic-antimycotic (cat no. 15240096; Thermo Fisher), and 20 mM HEPES at pH 7.4 for no more than 30 passages.

### Plasmids

LifeAct-mEmerald (no. 54148) and GCaMP6s (no. 40753) were obtained from Addgene (Watertown, MA). Eevee-ROCK was kindly provided by Dr. Michiyuki Matsuda (Kyoto University). 1 µg of plasmid was used to transfect 750,000 cells in each well of a 6-well plate using Lipofectamine 2000 (5 µL; Thermo Fisher) in OptiMEM (400 µL; Thermo Fisher). After 20 min at room temperature, plasmid in Lipofectamine 2000/OptiMEM was then incubated with cells in complete media (2 mL) overnight.

### Chemical treatments

BAPTA (cat no. 2786), GsMTx4 (Piezo1/2 inhibitor; cat no. 4912), and Yoda1 (Piezo1/2 activator; cat no. 5586) were purchased from Tocris Bioscience (Bristol, UK). Prior to confinement, cells were treated with drug for 1 hr. Simultaneously, confining devices were incubated with drug in complete media for at least 1 hr before loading cells. For measurements of intracellular [Ca^2+^], cells were simultaneously loaded with the red fluorescent Ca^2+^ indicator, Calbryte (cat no. 20700; AAT Bioquest, Pleasanton, CA), and a green fluorescent dye (cat no. C7025; Thermo Fisher) for ratiometric fluorescence imaging.

### RT-qPCR

Total RNA was isolated from cells using the PureLink RNA Mini Kit (cat no. 12183018A; Thermo Fisher) and was used for reverse transcription using a High-Capacity cDNA Reverse Transcription Kit (cat no. 4368814; Thermo Fisher). qPCR was performed using PowerUp SYBR Green Master Mix (cat no. A25742; Thermo Fisher) on a real-time PCR detection system (CFX96; Bio-Rad, Hercules, CA). Relative mRNA levels were calculated using the ΔCt method.

### RNA interference

Non-targeting (cat no. 4390844), Piezo1 (cat no. 4392420; s18891), cPLA2 (cat no. 4390824; s10592), MYLK (cat no. 4392420; s533772), ROCK1 (cat no. 4390824; s12097), ROCK2 (cat no. 4390824; s18161), MYPT1 (cat no. 4390824; s9235), and INF2 (cat no. 4392420; s230622) siRNAs were purchased from Thermo Fisher. All siRNA transfections were performed using RNAiMAX (5 µL; Thermo Fisher) and OptiMEM (400 µL; Thermo Fisher). 200,000 cells were trypsinized and seeded in 6-well plates in complete media. After cells adhered (∼1 hr), siRNAs in RNAiMAX/OptiMEM were added to cells in complete media (2 mL) at a final concentration of 50 nM. Cells were incubated with siRNAs for 3 days.

### 2D confinement

This protocol has been described in detail elsewhere [15]. Briefly, PDMS (cat no. 24236-10) was purchased from Electron Microscopy Sciences (Hatfield, PA). 2 mL was cured overnight at 37 °C in each well of a 6-well glass bottom plate (cat no. P06-1.5H-N; Cellvis, Mountain View, CA). Using a biopsy punch (cat no. 504535; World Precision Instruments, Sarasota, FL), an 8 mm hole was cut and 3 mL of serum free media containing 1% BSA was added to each well and incubated overnight at 37 °C. After removing the serum free media containing 1% BSA, 300 μL of complete media containing trypsinized cells (250,000 to 1 million) and 2 μL of 3.11 μm beads (cat no. PS05002; Bangs Laboratories, Fishers, IN) were then pipetted into the round opening.

The vacuum created by briefly lifting one side of the hole with a 1 mL pipette tip was used to move cells and beads underneath the PDMS. Finally, 3 mL of complete media was added to each well and cells were recovered for ∼60 min before imaging.

### Microchannel preparation

PDMS (cat no. 24236-10; Electron Microscopy Sciences) was prepared using a 1:7 ratio of base and curing agent. Uncured PDMS was poured over the wafer mold, placed in a vacuum chamber to remove bubbles, moved to a 37 °C incubator, and left to cure overnight. After curing, small PDMS slabs with microchannels were cut using a scalpel, whereas cell loading ports were cut using a 0.4 cm hole punch (cat no. 12-460-409; Fisher Scientific, Hampton, NH).

For making PDMS coated cover glass (cat no. 12-545-81; Fisher Scientific), 30 µL of uncured PDMS was pipetted at the center of the cover glass, placed in a modified mini-centrifuge, and spun for 30 sec for even spreading. The PDMS coated cover glass was then cured for at least 1 hr on a 95 °C hot plate.

Prior to slab and coated cover glass joining, PDMS surfaces were activated for ∼1 min by plasma treatment (cat no. PDC-32G; Harrick Plasma, Ithaca, NY). Immediately after activation, slabs were bonded to coated cover glass. For complete bonding, the apparatus was incubated at 37 °C for at least 1 hr.

### Microchannel coating

Prior to microchannel coating, surfaces were first activated by plasma treatment. VCAM-1 (cat no. 862-VC; R&D Systems, Minneapolis, MN) or BSA (cat no. VWRV0332; VWR, Radnor, PA) was used for coating at 1 µg/mL and 1%, respectively, in PBS. Immediately after plasma treatment, VCAM-1 or BSA solution was pumped into microchannels using a modified motorized pipette. To remove any bubbles pumped into microchannels, the apparatus was left to coat in a vacuum chamber for at least 1 hr. Afterward, VCAM-1 or BSA solution was aspirated out and microchannels were rinsed twice by pumping in PBS. Finally, microchannels were incubated in complete media over-night at 4 °C before use.

### Microchannel loading

Prior to cells being loaded into microchannels, complete media was aspirated, and microchannels were placed into an interchangeable cover-glass dish (cat no. 190310-35; Bioptechs, Butler, PA). Freshly trypsinized cells in 300 µL of complete media, stained with 1 µL far red membrane dye (cat no. C10046; Thermo Fisher), were pumped into microchannels using a modified motorized pipette. Once at least 20 cells are observed in microchannels by low magnification brightfield microscopy, microchannels were covered with 2 mL of complete media. Before imaging, a lid was placed on top the apparatus to prevent evaporation.

### Microscopy

High-resolution imaging was performed using a General Electric (Boston, MA) DeltaVision Elite imaging system mounted on an Olympus (Japan) IX71 stand with a motorized XYZ stage, Ultimate Focus, cage incubator, ultrafast solid-state illumination with excitation/emission filter sets for DAPI, CFP, FITC, GFP, YFP, TRITC, mCherry, and Cy5, critical illumination, Olympus PlanApo N 60X/1.42 NA DIC (oil) and UPlanSApo 60X/1.3 NA DIC (silicone) objectives, Photometrics (Tucson, AZ) CoolSNAP HQ2 camera, SoftWoRx software with constrained iterative deconvolution, and vibration isolation table.

### Cell classification in 2D confinement

Cells that displayed directionally persistent migration over at least 4 frames (32 min) were classified as leader mobile (LM). Any cell with a large bleb that remained stable for at least 4 frames (32 min) was considered to have a leader bleb.

### Cell classification in microchannels

Cells that moved at least ½ of its original length over a 5 hr timelapse movie were considered motile. Cells that displayed only blebs were classified as amoeboid, whereas cells with actin-based protrusions were classified as mesenchymal. Cells with blebs and actin-based protrusions were classified as hybrid.

### Cell morphology

Analyze_Blebs was used to measure the largest bleb area, aspect ratio, roundness, circularity, solidity, and Feret’s Diameter of cells from timelapse movies [16].

### Cell migration

To perform cell speed and directionality ratio analyses, we used an Excel (Microsoft, Redmond, WA) plugin, DiPer, developed by Gorelik and colleagues and the Fiji plugin, MTrackJ, developed by Erik Meijering for manual tracking [30]. Brightfield imaging was used to confirm that beads or debris were not obstructing the cell.

### FRET

Ratio images of FRET (CFP excitation/YFP emission) to CFP (CFP excitation/CFP emission) were generated and analyzed in Fiji (https://fiji.sc/).

### Cell Impedance

Cell impedance was measured using a Real Time Cell Analyzer (RTCA; Agilent, Santa Clara, CA). Before adding 30,000 cells to each well, plates (cat no. 300601140; Agilent) were coated with VCAM-1 (1 µg/mL) (cat no. 862-VC; R&D Systems) overnight at 4 °C.

### 2D confinement + adhesive micropatterns

Micropatterning was performed according to the manufacturer protocol (4Dcell, France). Briefly, coverslips were first coated with non-adhesive PLL-g-PEG. Lines were patterned on coverslips using a photomask (cat no. UM006; 4Dcell) and treatment with deep UV. The coverslips were then incubated with fibronectin (10 µg/ml) (cat no. 33016015; Thermo Fisher) for 1 hr at room temperature. To mark the adhesive patterns, a small amount of red fluorescent BSA (1 µg/mL) (cat no. A13101; Thermo Fisher) was added to the fibronectin solution. Cells were confined down to 3 µm using a dynamic cell confiner according to the manufacturer protocol (4Dcell). Because cells were found to be less motile in the dynamic cell confiner, all cells were treated with siRNA towards MYPT1 to generally increase rates of motility.

### Statistics

Sample sizes were determined empirically and based on saturation. As noted in each figure legend, statistical significance was determined by either a two-tailed Student’s t-test or multiple-comparison test post-hoc. Normality was determined by a D’Agostino & Pearson test in GraphPad Prism 7. * - p ≤ 0.05, ** - p ≤ 0.01, *** - p ≤ 0.001, and **** - p ≤ 0.0001

## Author Contributions

N.K.: Investigation, A.P.C.: Investigation, N.P.: Investigation, J.S.L.: Conceptualization, Supervision, Project administration, Visualization, Writing - Original Draft, Funding acquisition

## Acknowledgements

We would like to thank the Cady Lab (SUNY Polytechnic Institute, Albany, NY) for fabrication of silicon wafer molds.

---

**Figure 1-figure supplement 1.**
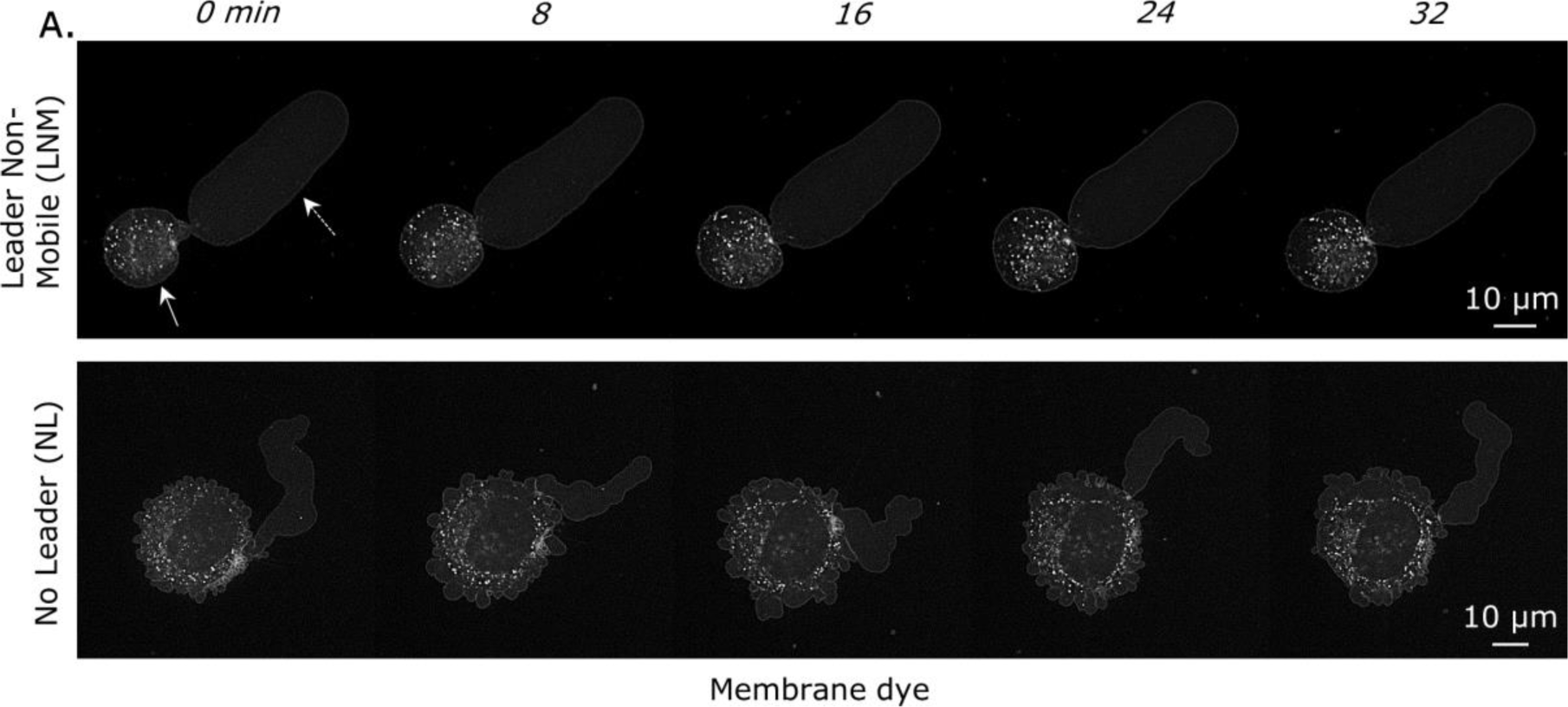
**A.** Montage of a leader non-mobile (LNM) and no leader (NL) cells vertically confined down to 3 µm. Cells were stained with a far red membrane dye. Arrows point to the cell body (solid) and leader bleb (dashed), which is a large and stable bleb. PDMS was coated with Bovine Serum Albumin (BSA; 1%).

**Figure 1-figure supplement 2.**
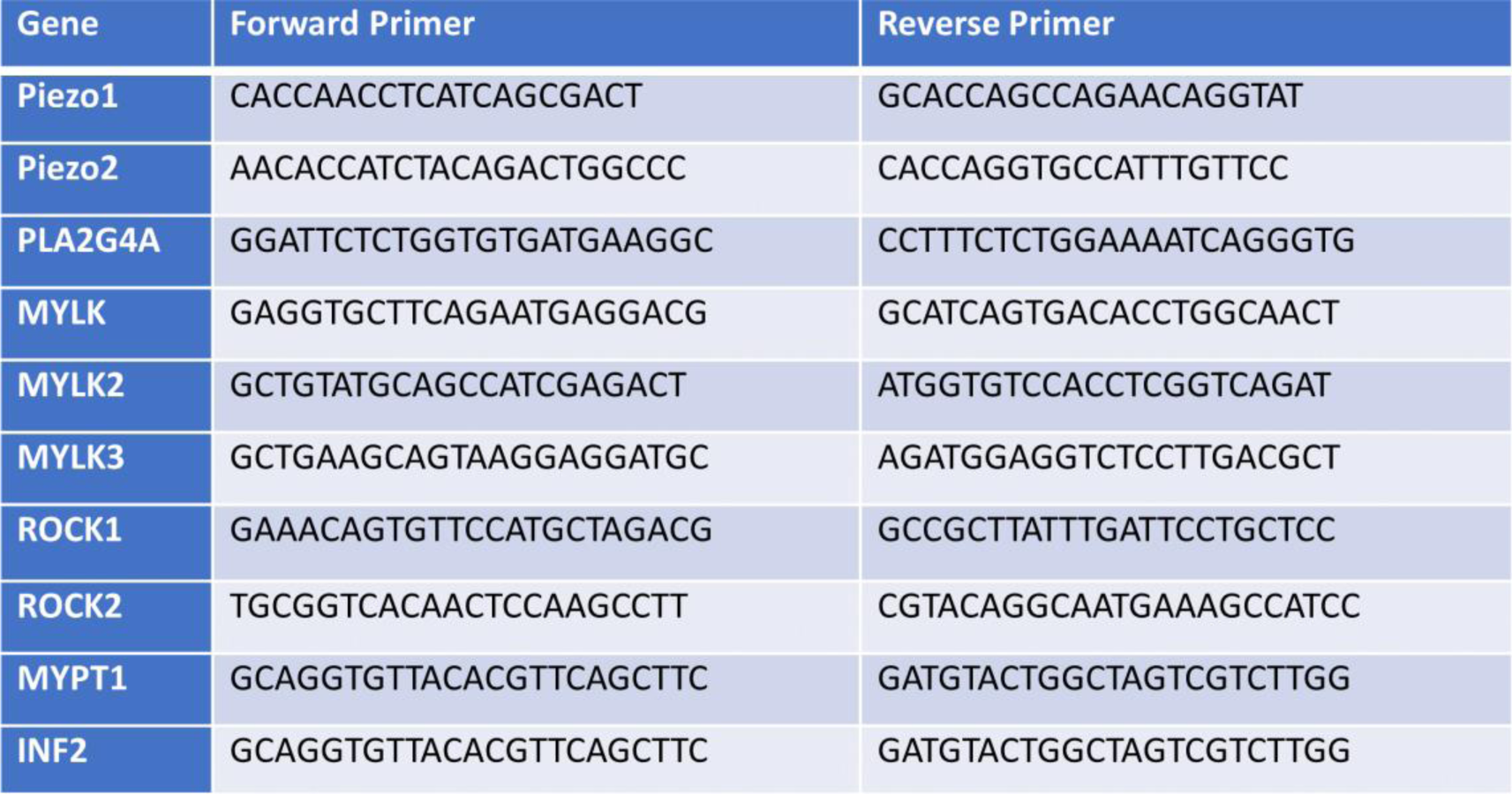
The listed qPCR primers were used to amplify Piezo1, Piezo2, cPLA2 (PLA2G4A), MYLK., MYLK2, MYLK3, ROCK1, ROCK2, MYPT1, and INF2 from cDNA.

**Figure 1-figure supplement 3.**
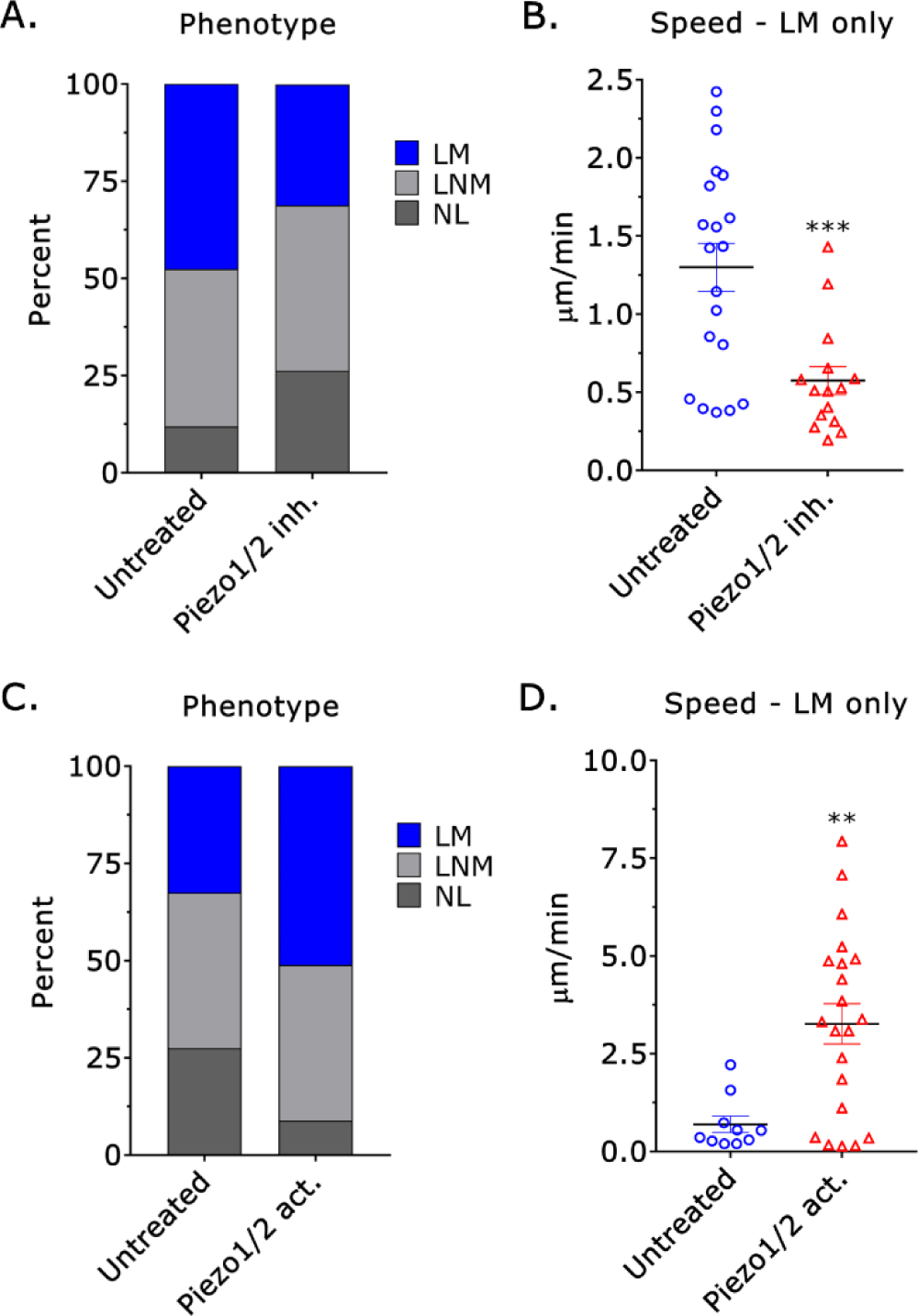
**A.** Compared to untreated (n=42), fewer leader mobile (LM) cells were observed after treatment with a Piezo1/2 inhibitor (GsMTx4; 10 µM) (n=61) (χ^2^ ≤ 0.0001). **B.** Compared to untreated (n=20), Piezo1/2 inhibitor (GsMTx4; 10 µM) (n=18) treated leader mobile (LM) cells were significantly slower (Student’s t-test; mean +/- SEM). **C.** Compared to untreated (n=40), more leader mobile (LM) cells were observed after treatment with a Piezo1/2 activator (Yoda1; 20 µM) (n=45; χ^2^ ≤ 0.0001). **D.** Compared to untreated (n=10), Piezo1/2 activator (Yoda1; 20 µM) (n=21) treated leader mobile (LM) cells were significantly faster (Student’s t-test; mean +/- SEM). PDMS was coated with BSA (1%). * - p ≤ 0.05, ** - p ≤ 0.01, *** - p ≤ 0.001, and **** - p ≤ 0.0001

**Figure 1-figure supplement 4.**
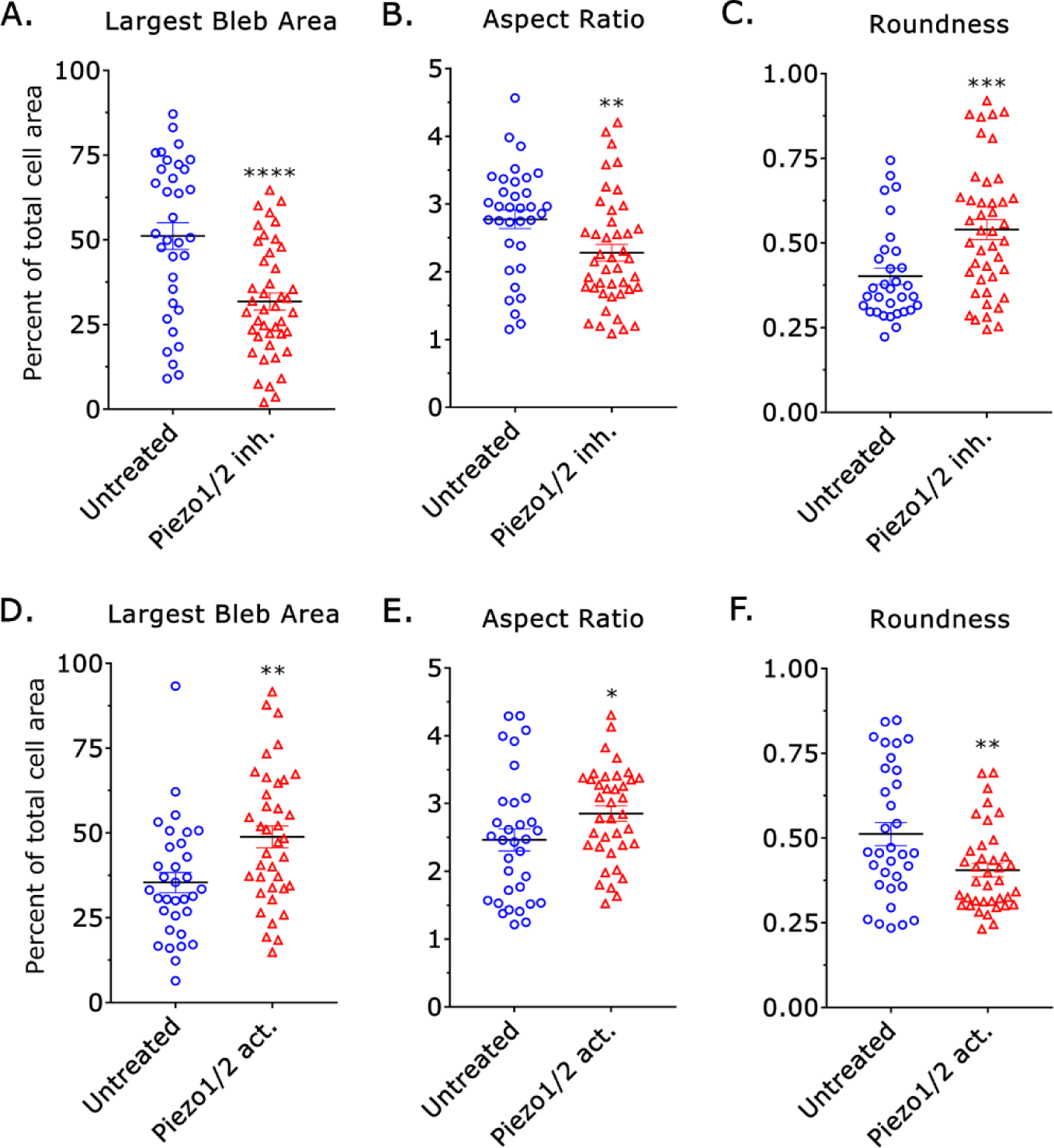
**A.-C.** Compared to untreated (n=34), the largest bleb area, aspect ratio, and roundness of cells were significantly different after Piezo1 inhibitor (GsMTx4; 10 µM) (n=43) treatment (Student’s t-test; mean +/- SEM). **D.-F.** Compared to untreated (n=33), the largest bleb area, aspect ratio, and roundness of cells were significantly different after Piezo1 activator (Yoda1; 20 µM) (n=38) treatment (Student’s t-test; mean +/- SEM). Analyze_Blebs was used to measure shape descriptors. PDMS was coated with BSA (1%). * - p ≤ 0.05, ** - p ≤ 0.01, *** - p ≤ 0.001, and **** - p ≤ 0.0001

**Figure 4-figure supplement 1.**
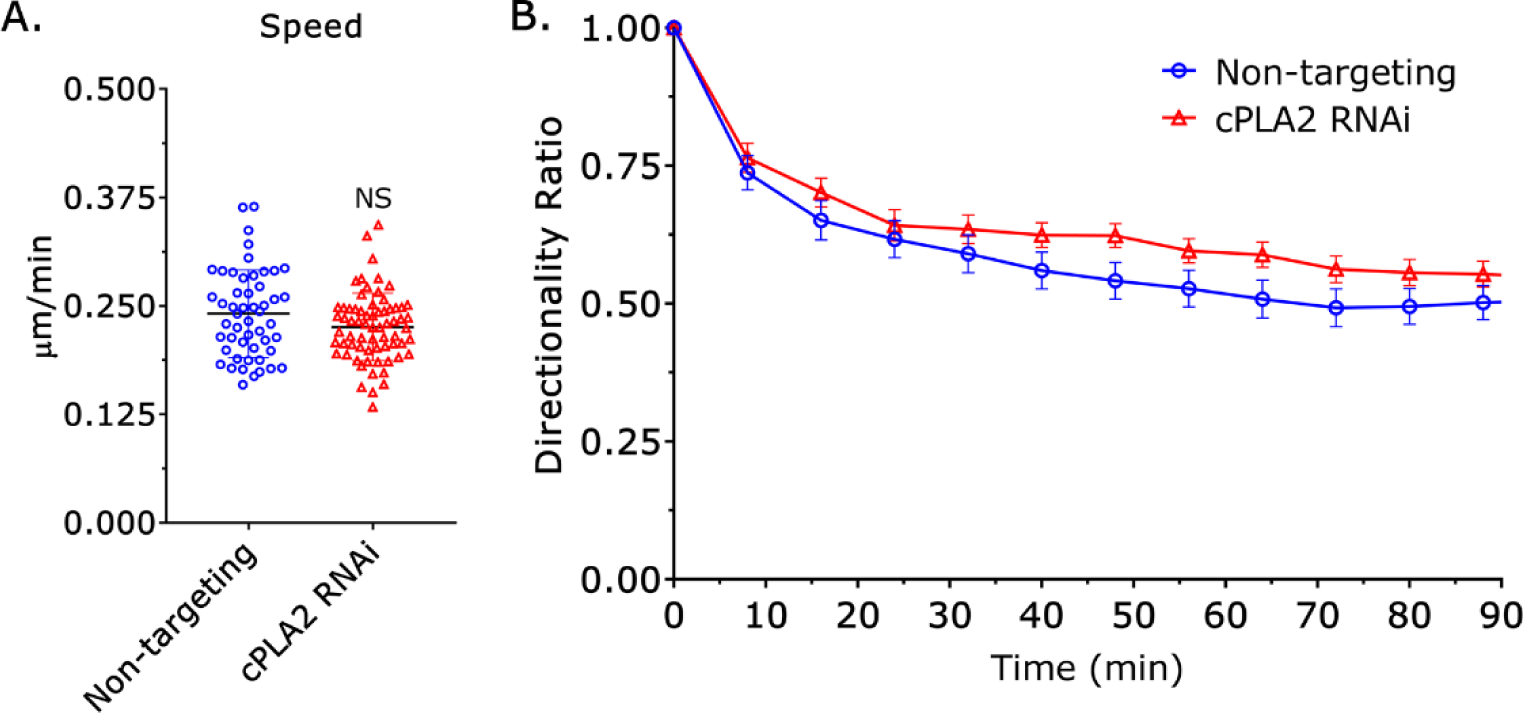
**A.** Based on manual cell tracking, the speed of non-targeting and cPLA2 siRNA treated cells were similar (Student’s t-test; mean +/- SEM). **B.** Based on manual cell tracking, the directionality ratio for non-targeting (n=51) and cPLA2 siRNA (n=69) treated cells were similar (mean +/- SEM). Microchannels are coated with VCAM-1 (1 µg/mL) and are 8 µm in height, 8 µm in width, and 100 µm in length.

**Figure 5-figure supplement 1.**
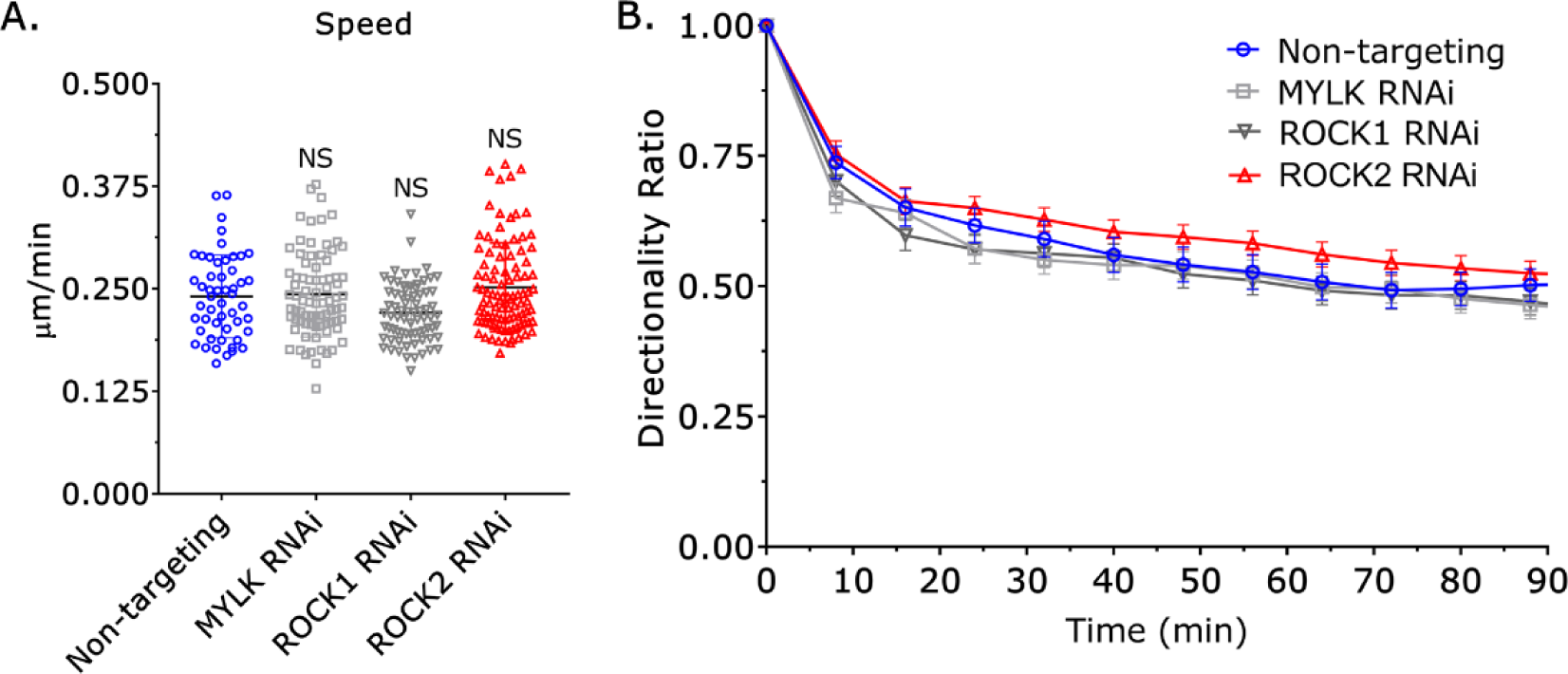
**A.** Based on manual cell tracking, the speed of non-targeting, MYLK, ROCK1, and ROCK2 siRNA treated cells were similar (Student’s t-test; mean +/- SEM). **B.** Based on manual cell tracking, the directionality ratio for non-targeting (n=51), MYLK (n=84), ROCK1 (n=75), and ROCK2 (n=100) siRNA treated cells were similar (mean +/- SEM). Microchannels are coated with VCAM-1 (1 µg/mL) and are 8 µm in height, 8 µm in width, and 100 µm in length. Microchannels are coated with VCAM-1 (1 µg/mL) and are 8 µm in height, 8 µm in width, and 100 µm in length.

**Figure 5-figure supplement 2.**
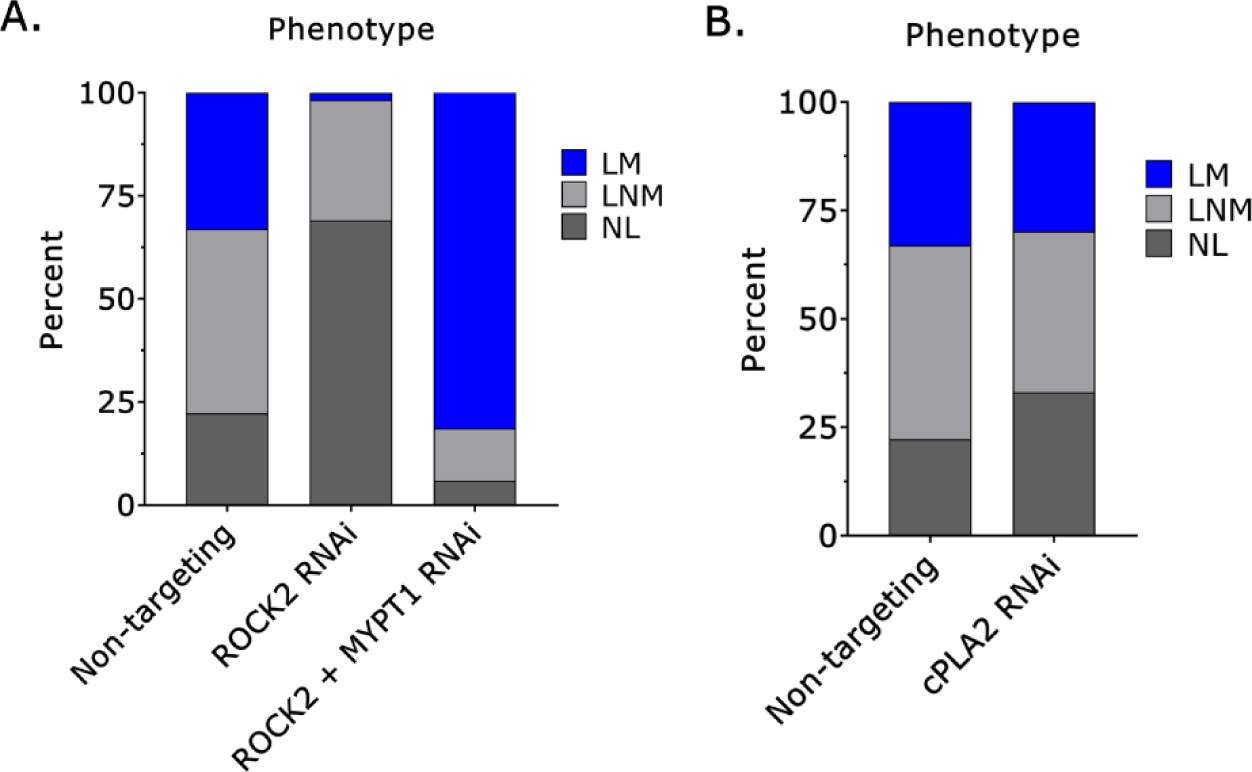
**A.** Percent leader mobile (LM), leader non-mobile (LNM), and no leader (NL) cells treated with a non-targeting (n=57), ROCK2 (n=67; χ^2^ ≤ 0.0001), and with both a ROCK2 and MYPT1 siRNA (n=118; χ^2^ ≤ 0.0001). **B.** Percent leader mobile (LM), leader non-mobile (LNM), and no leader (NL) cells treated with a non-targeting (n=57) or cPLA2 siRNA (n=82; χ^2^ ≤ 0.05). PDMS was coated with BSA (1%).

**Figure 5-figure supplement 3.**
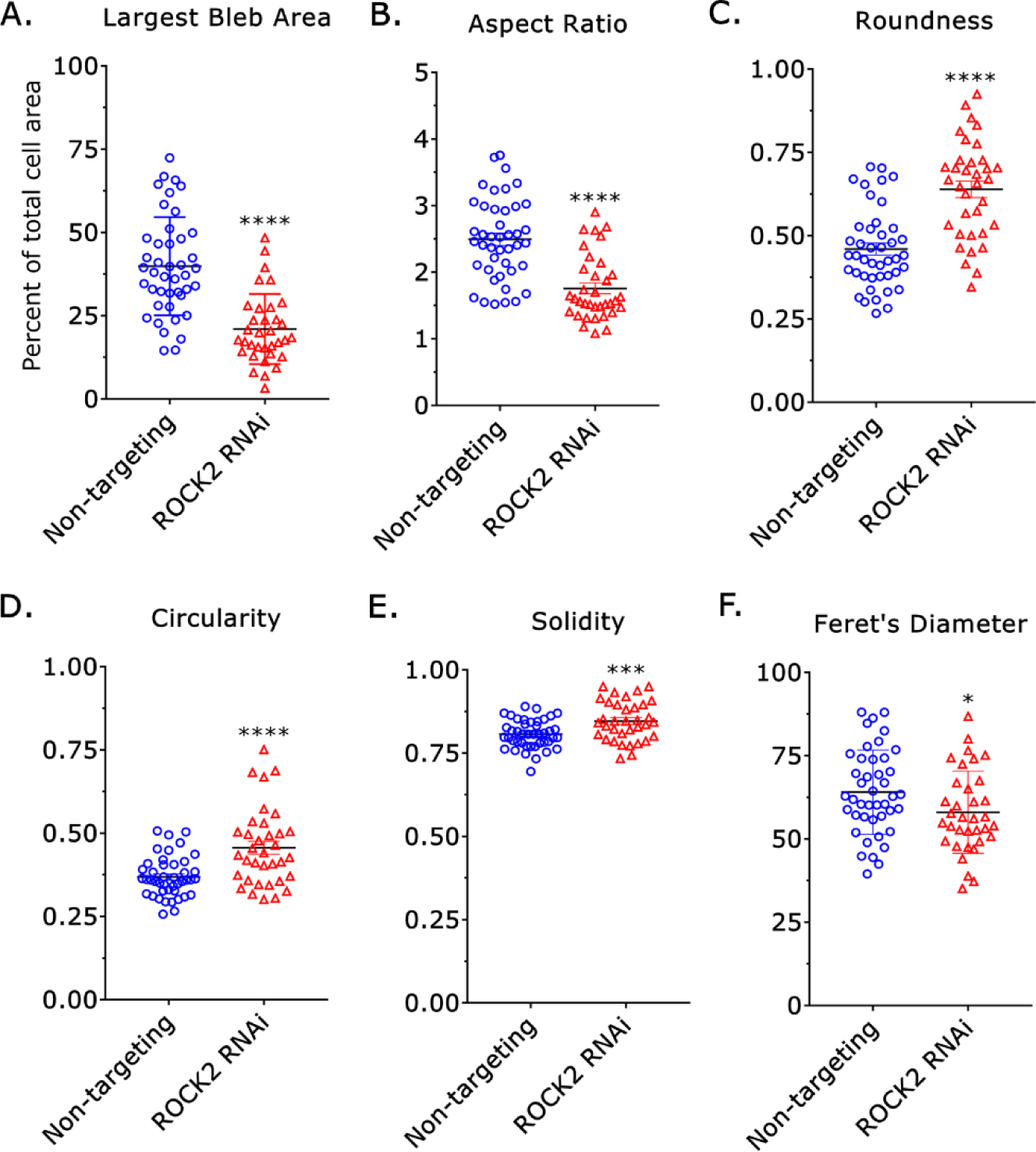
**A.-F.** Compared to non-targeting (n=43), the largest bleb area, aspect ratio, roundness, circularity, solidity, and Feret’s Diameter of cells were significantly different after treatment with a ROCK2 siRNA (n=35) (Student’s t-test; mean +/- SEM). Analyze_Blebs was used to measure shape descriptors. PDMS was coated with BSA (1%). * - p ≤ 0.05, ** - p ≤ 0.01, *** - p ≤ 0.001, and **** - p ≤ 0.0001

**Figure 6-figure supplement 1.**
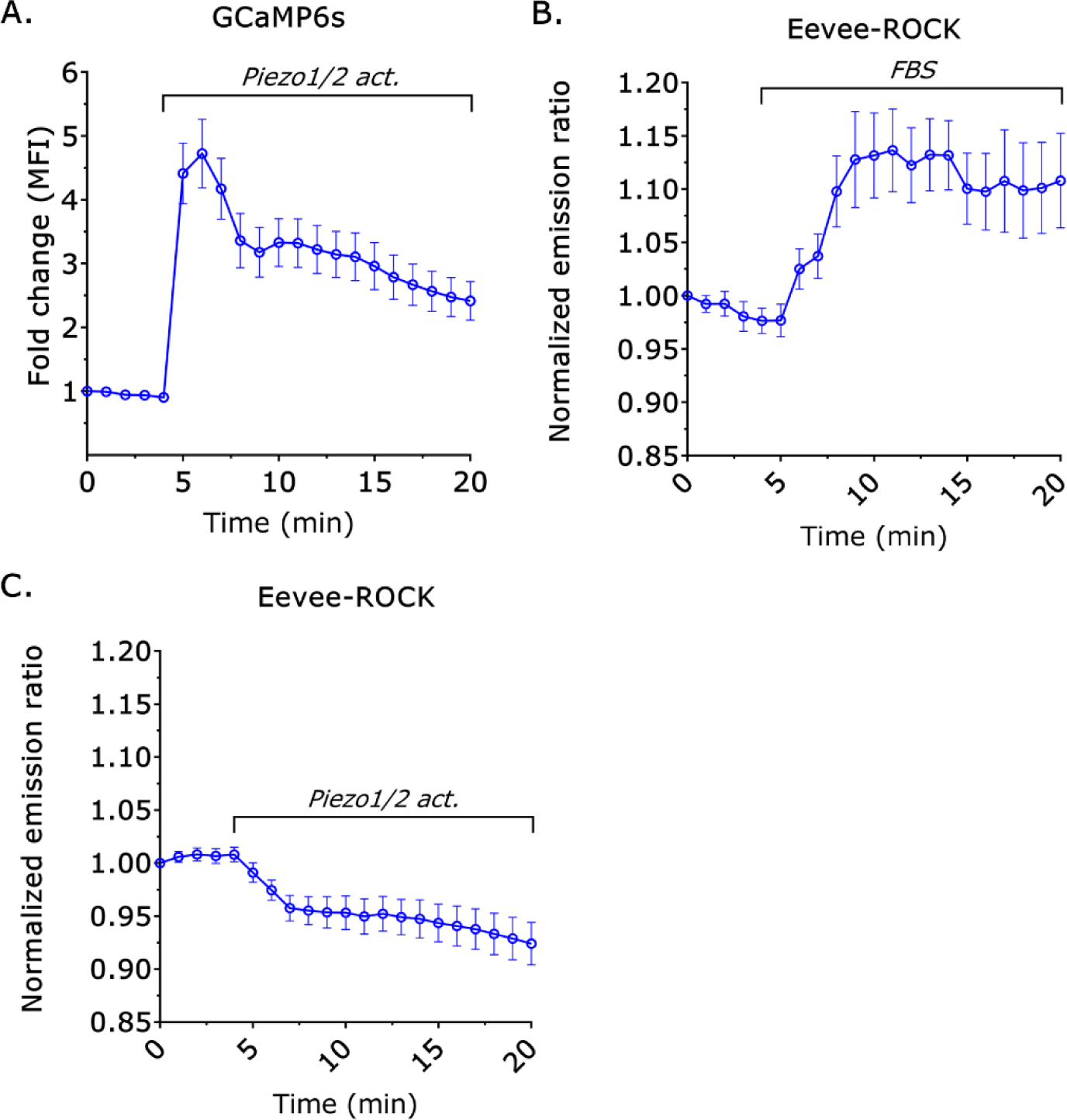
**A.** Fold change in mean fluorescence intensity (MFI) for GCaMP6s over time (mean +/- SEM). Piezo1/2 activator (Yoda1; 10 µM) was added to cells after 5 min (n=19). **B.** Normalized emission ratio or Fluorescence Resonance Energy Transfer (FRET) for Eevee-ROCK over time (mean +/- SEM). Fetal Bovine Serum (FBS; 10%) was added after 5 min (n=9). **C.** Normalized emission ratio or FRET for Eevee-ROCK over time (mean +/- SEM). Piezo1/2 activator (Yoda1; 10 µM) was added to cells after 5 min (n=21). Cells were plated on cover glass.

**Figure 6-figure supplement 2.**
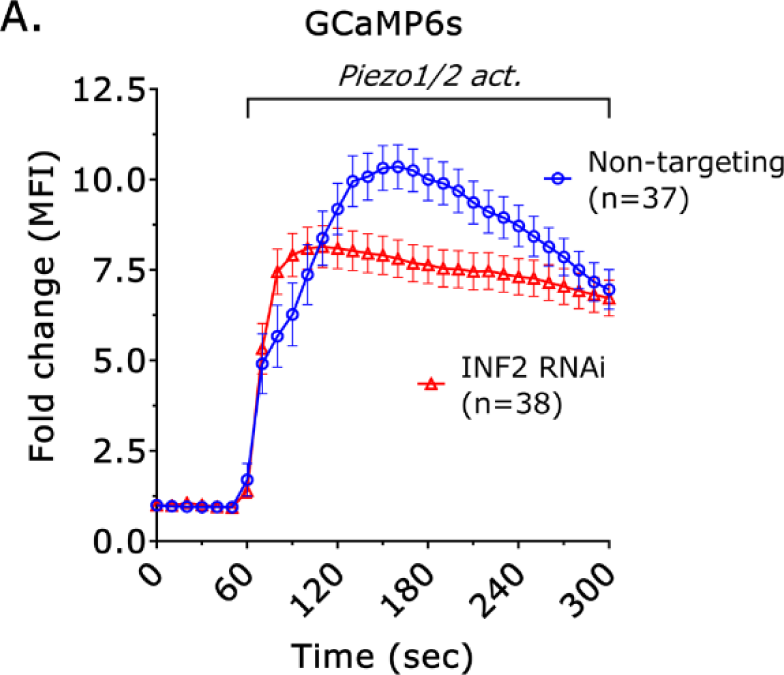
**A.** Fold change in mean fluorescence intensity (MFI) for GCaMP6s over time (mean +/- SEM). Piezo1/2 activator (Yoda1; 10 µM) was added to cells after 60 sec. Cells were treated with a non-targeting or INF2 siRNA and plated on cover glass.

**Figure 7-figure supplement 1.**
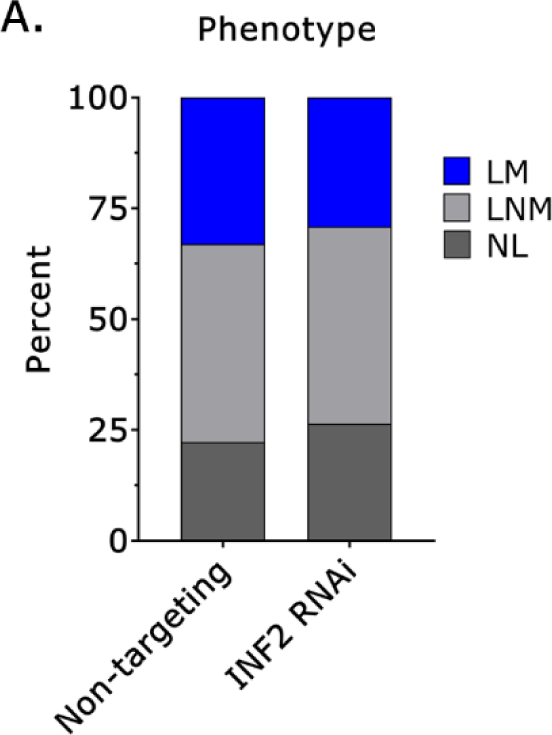
**A.** Percent leader mobile (LM), leader non-mobile (LNM), and no leader (NL) cells treated with a non-targeting (n=57) or INF2 siRNA (n=98; χ^2^=NS). PDMS was coated with BSA (1%).

**Figure 8-figure supplement 1.**
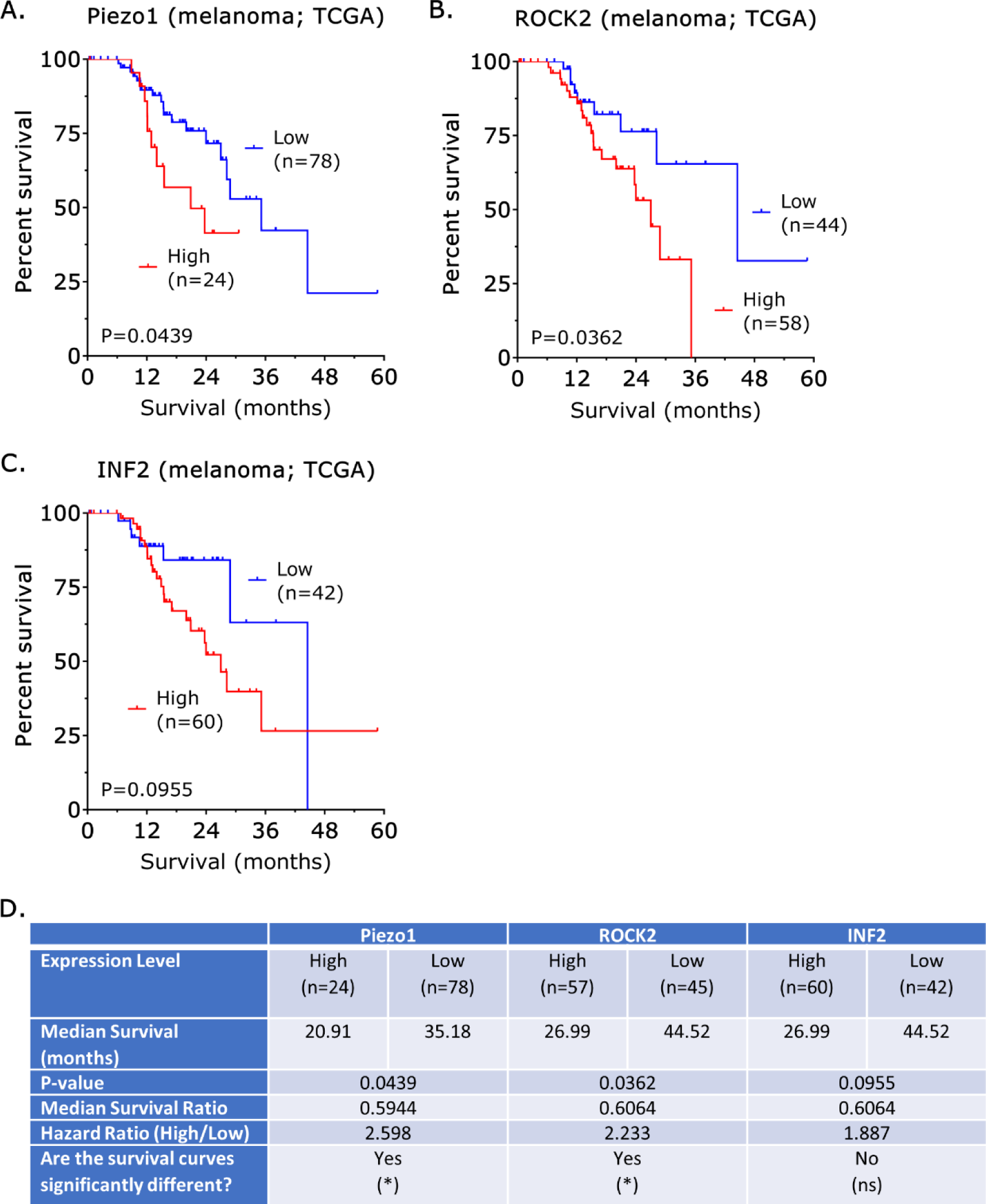
**A.-C.** Survival curves for patients with high vs. low Piezo1, ROCK2, and INF2 mRNA levels. **D.** Table summarizing survival curves for Piezo1, ROCK2, and INF2. All survival curves are based on RNA sequencing data from The Cancer Genome Atlas (TCGA; https://www.cancer.gov/tcga) for melanoma. A cut-off value yielding the lowest p-value on a log-rank test was used in each. * - p ≤ 0.05, ** - p ≤ 0.01, *** - p ≤ 0.001, and **** - p ≤ 0.0001

## Key Resources Table

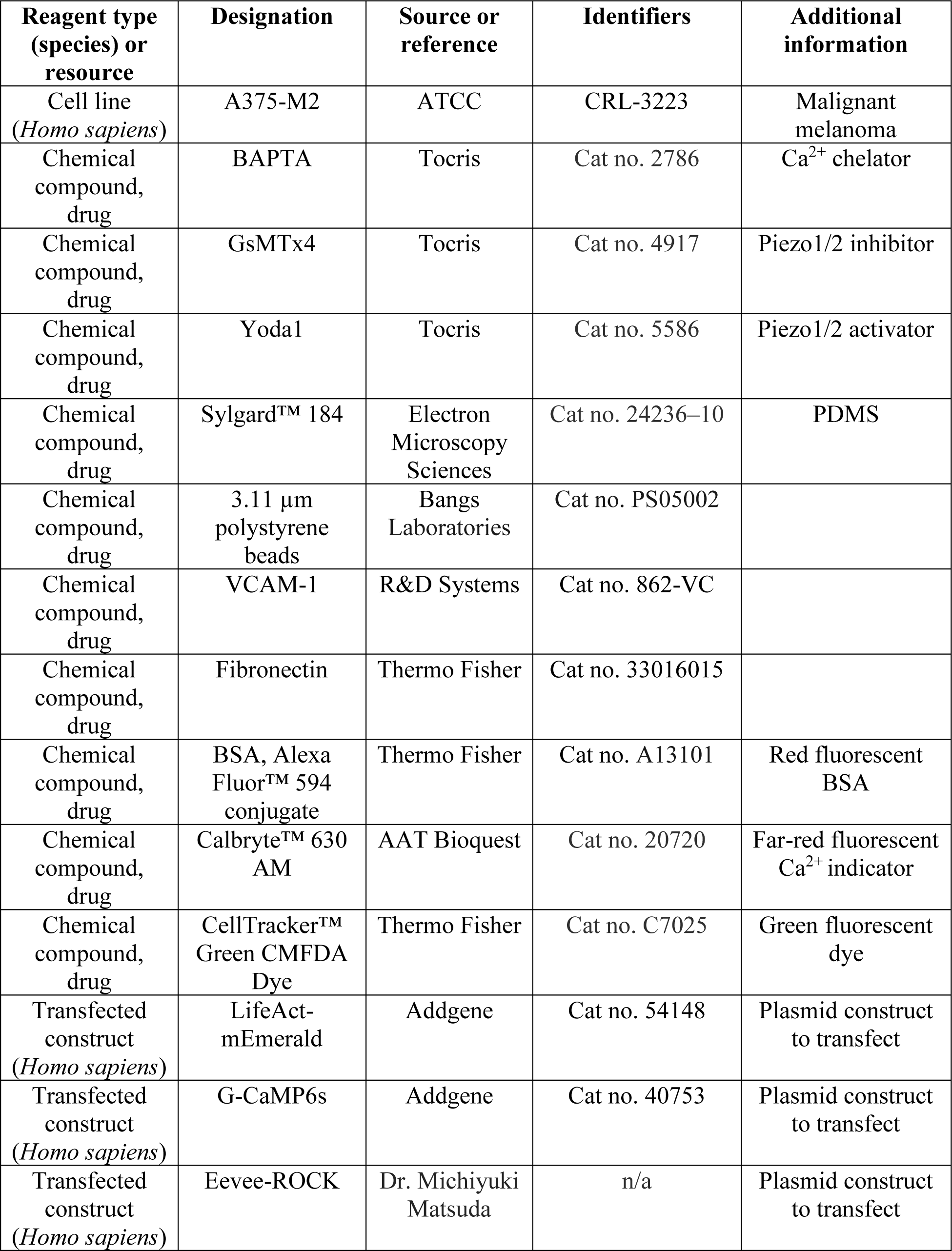
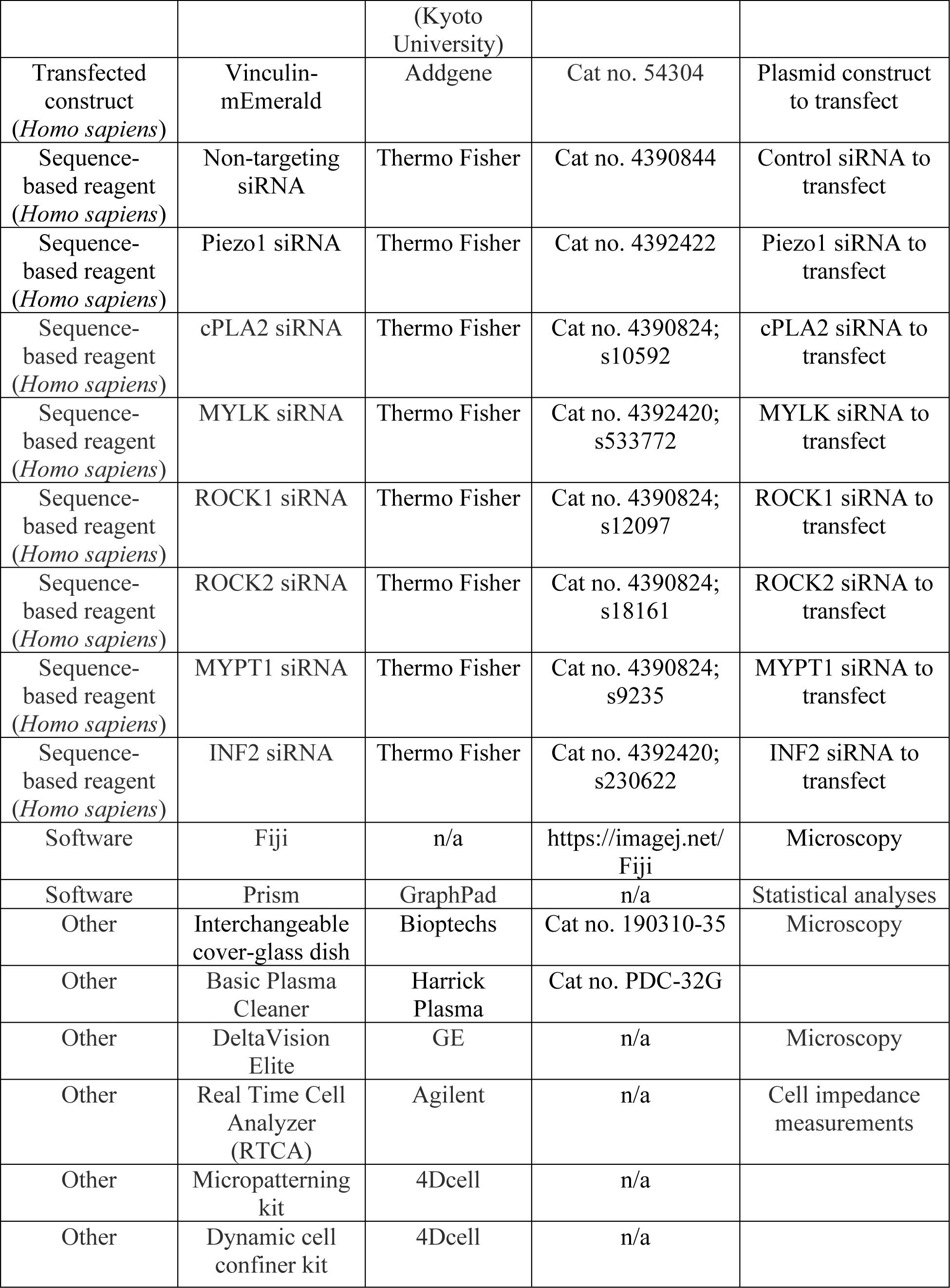

## Videos

**Figure 1-video 1.** A leader mobile (LM) cell confined down to 3 µm. Cells were stained with a far red membrane dye. PDMS was coated with Bovine Serum Albumin (BSA; 1%). Corresponds to Figure 1A.

**Figure 1-video 2.** A leader non- mobile (LNM) cell confined down to 3 µm. Cells were stained with a far red membrane dye. PDMS was coated with BSA (1%). Corresponds to Figure 1-figure supplement 1A.

**Figure 1-video 3.** A no leader (NL) cell confined down to 3 µm. Cells were stained with a far red membrane dye. PDMS was coated with BSA (1%). Corresponds to Figure 1-figure supplement 1A.

**Figure 2-video 1.** A motile cell adopting a hybrid phenotype. Cells were stained with a far red membrane dye. Microchannels are coated with VCAM-1 (1 µg/mL) and are 8 µm in height, 8 µm in width, and 100 µm in length. Corresponds to Figure 2A.

**Figure 2-video 2.** A motile cell adopting an amoeboid phenotype in an undulating channel. Cells were stained with a far red membrane dye. Microchannels are coated with VCAM-1 (1 µg/mL) and are 8 µm in height, 8 µm in width, and 100 µm in length. Corresponds to Figure 2C.

**Figure 6-video 1.** Cell transiently transfected with a non-targeting siRNA and LifeAct- mEmerald treated with a Piezo1/2 activator (Yoda1; 20 µM). Cell was plated on fibronectin coated (10 µg/mL) glass. Corresponds to Figure 6B.

**Figure 6-video 2.** Cell transiently transfected with an INF2 siRNA and LifeAct-mEmerald treated with a Piezo1/2 activator (Yoda1; 20 µM). Cell was plated on fibronectin coated (10 µg/mL) glass. Corresponds to Figure 6B.

**Figure 7-video 1.** An INF2 siRNA treated cell adopting a mesenchymal migration mode in an undulating channel. Cells were stained with a far red membrane dye. Microchannels are coated with VCAM-1 (1 µg/mL) and are 8 µm in height, 8 µm in width, and 100 µm in length. Corresponds to Figure 7A.

**Figure 8-video 1.** A non-targeting siRNA treated cell (green) migrating across an adhesive (fibronectin; 10 µg/mL) micropattern (magenta). The cell was stained with a green fluorescent dye. Cells were confined down to 3 µm using a dynamic cell confiner (4Dcell). Corresponds to Figure 8C.

**Figure 8-video 2.** INF2 siRNA treated cells (green) adhere to the adhesive (fibronectin; 10 µg/mL) micropattern (magenta). The cells were stained with a green fluorescent dye. Cells were confined down to 3 µm using a dynamic cell confiner (4Dcell). Corresponds to Figure 8D.

## References

1. Paul, C.D., Mistriotis, P., and Konstantopoulos, K. (2017). Cancer cell motility: lessons from migration in confined spaces. Nature Reviews Cancer 17, 131.

2. Liu, Y.J., Le Berre, M., Lautenschlaeger, F., Maiuri, P., Callan-Jones, A., Heuze, M., Takaki, T., Voituriez, R., and Piel, M. (2015). Confinement and low adhesion induce fast amoeboid migration of slow mesenchymal cells. Cell 160, 659–672.

3. Ruprecht, V., Wieser, S., Callan-Jones, A., Smutny, M., Morita, H., Sako, K., Barone, V., Ritsch-Marte, M., Sixt, M., Voituriez, R., et al. (2015). Cortical contractility triggers a stochastic switch to fast amoeboid cell motility. Cell 160, 673–685.

4. Logue, J.S., Cartagena-Rivera, A.X., Baird, M.A., Davidson, M.W., Chadwick, R.S., and Waterman, C.M. (2015). Erk regulation of actin capping and bundling by Eps8 promotes cortex tension and leader bleb-based migration. Elife 4.

5. Yamada, K.M., and Sixt, M. (2019). Mechanisms of 3D cell migration. Nature Reviews Molecular Cell Biology 20, 738–752.

6. Charras, G.T., Yarrow, J.C., Horton, M.A., Mahadevan, L., and Mitchison, T.J. (2005). Non-equilibration of hydrostatic pressure in blebbing cells. Nature 435, 365–369.

7. Charras, G.T., Coughlin, M., Mitchison, T.J., and Mahadevan, L. (2008). Life and times of a cellular bleb. Biophys J 94, 1836–1853.

8. Tozluoglu, M., Tournier, A.L., Jenkins, R.P., Hooper, S., Bates, P.A., and Sahai, E. (2013). Matrix geometry determines optimal cancer cell migration strategy and modulates response to interventions. Nat Cell Biol 15, 751–762.

9. Bergert, M., Erzberger, A., Desai, R.A., Aspalter, I.M., Oates, A.C., Charras, G., Salbreux, G., and Paluch, E.K. (2015). Force transmission during adhesion-independent migration. Nat Cell Biol 17, 524–529.

10. Hung, W.C., Yang, J.R., Yankaskas, C.L., Wong, B.S., Wu, P.H., Pardo-Pastor, C., Serra, S.A., Chiang, M.J., Gu, Z., Wirtz, D., et al. (2016). Confinement Sensing and Signal Optimization via Piezo1/PKA and Myosin II Pathways. Cell Rep 15, 1430–1441.

11. Lomakin, A.J., Cattin, C.J., Cuvelier, D., Alraies, Z., Molina, M., Nader, G.P.F., Srivastava, N., Saez, P.J., Garcia-Arcos, J.M., Zhitnyak, I.Y., et al. (2020). The nucleus acts as a ruler tailoring cell responses to spatial constraints. Science 370.

12. Venturini, V., Pezzano, F., Catala Castro, F., Hakkinen, H.M., Jimenez-Delgado, S., Colomer-Rosell, M., Marro, M., Tolosa-Ramon, Q., Paz-Lopez, S., Valverde, M.A., et al. (2020). The nucleus measures shape changes for cellular proprioception to control dynamic cell behavior. Science 370.

13. Enyedi, B., Jelcic, M., and Niethammer, P. (2016). The Cell Nucleus Serves as a Mechanotransducer of Tissue Damage-Induced Inflammation. Cell 165, 1160–1170.

14. Shen, Z., Belcheva, K.T., Jelcic, M., Hui, K.L., Katikaneni, A., and Niethammer, P. (2022). A synergy between mechanosensitive calcium- and membrane-binding mediates tension-sensing by C2-like domains. Proc Natl Acad Sci U S A 119.

15. Jeremy, L., Clare, W., and Richard, C. (2018). A simple method for precisely controlling the confinement of cells in culture. In Protocol Exchange.

16. Vosatka, K.W., Lavenus, S.B., and Logue, J.S. (2022). A novel Fiji/ImageJ plugin for the rapid analysis of blebbing cells. PLoS One 17, e0267740.

17. Gabbireddy, S.R., Vosatka, K.W., Chung, A.J., and Logue, J.S. (2021). Melanoma cells adopt features of both mesenchymal and amoeboid migration within confining channels. Sci Rep 11, 17804.

18. Kimura, K., Ito, M., Amano, M., Chihara, K., Fukata, Y., Nakafuku, M., Yamamori, B., Feng, J., Nakano, T., Okawa, K., et al. (1996). Regulation of myosin phosphatase by Rho and Rho-associated kinase (Rho-kinase). Science 273, 245–248.

19. Li, C., Imanishi, A., Komatsu, N., Terai, K., Amano, M., Kaibuchi, K., and Matsuda, M. (2017). A FRET Biosensor for ROCK Based on a Consensus Substrate Sequence Identified by KISS Technology. Cell Struct Funct 42, 1–13.

20. Wales, P., Schuberth, C.E., Aufschnaiter, R., Fels, J., Garcia-Aguilar, I., Janning, A., Dlugos, C.P., Schafer-Herte, M., Klingner, C., Walte, M., et al. (2016). Calcium-mediated actin reset (CaAR) mediates acute cell adaptations. Elife 5.

21. Chhabra, E.S., Ramabhadran, V., Gerber, S.A., and Higgs, H.N. (2009). INF2 is an endoplasmic reticulum-associated formin protein. J Cell Sci 122, 1430–1440.

22. Riedl, J., Crevenna, A.H., Kessenbrock, K., Yu, J.H., Neukirchen, D., Bista, M., Bradke, F., Jenne, D., Holak, T.A., Werb, Z., et al. (2008). Lifeact: a versatile marker to visualize F-actin. Nat Methods 5, 605–607.

23. George, S., Martin, J.A.J., Graziani, V., and Sanz-Moreno, V. (2022). Amoeboid migration in health and disease: Immune responses versus cancer dissemination. Front Cell Dev Biol 10, 1091801.

24. Pandya, P., Orgaz, J.L., and Sanz-Moreno, V. (2017). Modes of invasion during tumour dissemination. Mol Oncol 11, 5–27.

25. Sanz-Moreno, V., Gadea, G., Ahn, J., Paterson, H., Marra, P., Pinner, S., Sahai, E., and Marshall, C.J. (2008). Rac activation and inactivation control plasticity of tumor cell movement. Cell 135, 510–523.

26. Lavenus, S.B., Vosatka, K.W., Caruso, A.P., Ullo, M.F., Khan, A., and Logue, J.S. (2022). Emerin regulation of nuclear stiffness is required for fast amoeboid migration in confined environments. J Cell Sci 135.

27. Sanz-Moreno, V., Gaggioli, C., Yeo, M., Albrengues, J., Wallberg, F., Viros, A., Hooper, S., Mitter, R., Feral, C.C., Cook, M., et al. (2011). ROCK and JAK1 signaling cooperate to control actomyosin contractility in tumor cells and stroma. Cancer Cell 20, 229–245.

28. Shoji, K.F., Bayet, E., Leverrier-Penna, S., Le Devedec, D., Mallavialle, A., Marionneau-Lambot, S., Rambow, F., Perret, R., Joussaume, A., Viel, R., et al. (2023). The mechanosensitive TRPV2 calcium channel promotes human melanoma invasiveness and metastatic potential. EMBO Rep, e55069.

29. Kamm, K.E., and Stull, J.T. (2001). Dedicated myosin light chain kinases with diverse cellular functions. J Biol Chem 276, 4527–4530.

30. Meijering, E., Dzyubachyk, O., and Smal, I. (2012). Methods for cell and particle tracking. Methods Enzymol 504, 183–200.

